# DDX3 Regulates the Innate Immune Response to Bone Sarcomas

**DOI:** 10.64898/2026.07.01.735844

**Authors:** Rachel Weil, Esteban Uceda Arias-Stella, Da Peng, Patrick Cahan, Natalie ter Hoeve, Paul J. van Diest, Venu Raman, Poornima Gourabathini, Kimberly Q. McKinney, Kenzie Wells, Kaitlyn H. Smith, Jeffrey Huo, Javier Oesterheld, David M. Loeb

## Abstract

Osteosarcoma (OS) and Ewing sarcoma (EWS) are the most common malignant bone tumors in children and adolescents, with survival rates around 25% in metastatic disease and few advances in treatment in decades. High DDX3 expression has been reported across various sarcoma subtypes. Depending on the context, DDX3 appears to have opposing roles in regulating the tumor immune microenvironment. Within macrophages, DDX3 promotes inflammatory cytokine expression and supports immune cell function. In contrast, in tumor cells DDX3 suppresses a pro-inflammatory state by unwinding dsRNAs, preventing a Type I interferon response. We show that inhibiting DDX3 with RK-33 leads to dsRNA accumulation, inducing a Type I interferon response and broader inflammatory gene expression changes across multiple sarcoma models, shifting macrophage polarization toward a pro-inflammatory M1-like phenotype. To evaluate whether this innate immune microenvironmental remodeling could translate into clinical benefit, we assessed the therapeutic efficacy of RK-33 alone or in combination with mifamurtide, an immunostimulant, in immune competent mouse models of osteosarcoma, with metastatic burden as the primary outcome. We found that in the absence of MYC over-expression, the combination treatment significantly reduced metastatic spread. These findings support targeting DDX3 as a novel innate immune based therapeutic strategy and highlight that the tumor’s molecular landscape critically influences therapeutic responsiveness.

## Introduction

Osteosarcoma (OS) and Ewing sarcoma (EWS) are the most common malignant bone tumors of children, adolescents, and young adults^1^. Patients diagnosed with metastatic or relapsed disease face poor survival outcomes, with few effective treatment options available. Immunotherapies show broad promise across a number of cancer types, but to date have achieved limited success in sarcomas. While many immunotherapies, including checkpoint inhibitors^2^, focus on the adaptive immune response, the innate immune system has received significantly less attention. The innate immune system serves as the first line of defense in anti-tumor immunity by directly recognizing and eliminating tumor cells^3^. In addition, innate immune cells regulate adaptive immune responses and retain capacity to exert anti-tumor effects even within an immunosuppressive environment^4^. Targeting innate immune pathways can effectively recruit and activate components of the adaptive immune system, thereby weakening tumor immune evasion mechanisms^5^. Macrophages are a key cellular mediator of this process, serving as a vital link between innate and adaptive immunity through functions such as antigen presentation and secretion of cytokines^6^. Macrophage behavior is highly context dependent, and macrophages can display opposite functions based on cues from the microenvironment^7^. Tumor associated macrophages (TAMs) often contribute to the immunosuppressive tumor microenvironment (TME), driving growth, metastasis and drug resistance^8,9^. Depending on their environment, macrophages undergo polarization between M1 type macrophages, associated with anti-tumoral, pro-inflammatory activity, and M2 type macrophages which are anti-inflammatory and promote tumorigenesis^6,10^. Altering the makeup of the TME can polarize macrophages toward an M1-like phenotype, harnessing their inflammatory power to actively assist in tumor clearance.

DDX3 is an RNA helicase upregulated in many cancers^11,12,13,14,15,16^, including sarcomas^17^. Aside from its roles in RNA metabolism^18,19^, DDX3 plays a role in the function of the innate immune system, promoting the production of inflammatory cytokines that modulate immune cell function. TANK-binding kinase 1 (TBK1) phosphorylates DDX3, and phosphorylated DDX3 interacts with IFN-α/β promoters leading to the transcription of cytokines, chemokines, and type-I interferons^20^ which activate a protective immune response. DDX3 can influence the nuclear factor kappa B (NF-kB) signaling pathway as well, affecting production of various inflammatory cytokines (including IL-12 and IFN-γ)^21^. In macrophages, DDX3 interacts with NLRP3 to drive inflammasome activation, leading to a caspase-1 dependent release of pro-inflammatory cytokines and pyroptotic cell death^22^. Although within cells of the innate immune system DDX3 drives a pro-inflammatory response^18^, when expressed in cancer cells, DDX3 may be anti-inflammatory. Studies in breast cancer found that DDX3 can unwind double stranded RNAs (dsRNAs), thereby preventing the activation of an inflammatory response^23^. Upon inhibition of DDX3, dsRNAs accumulate in the cytoplasm of cancer cells, triggering a downstream pathway leading to a Type I interferon response^23^. RK-33^24,25^, a small molecule inhibitor of DDX3, is selectively cytotoxic to sarcoma cell lines *in vitro* and slows the growth of high DDX3 expressing patient-derived xenograft (PDX) tumors growing in immune deficient mice^17^. Because DDX3 plays a pro-inflammatory role in immune cells but an anti-inflammatory role in cancer cells, it is not clear, *a priori*, whether inhibiting DDX3 will have an overall pro- or anti-inflammatory effect when treating an animal with a tumor. Because cancer cells far outnumber macrophages in the tumor microenvironment, we hypothesized that RK-33, by inhibiting DDX3, promotes overall pro-inflammatory changes. Our findings, both *in vitro* and *in vivo*, show that inhibiting DDX3 in cancer cells and in intact tumors leads to activation of a Type I interferon response, induces a significant pro-inflammatory polarization of macrophages in multiple *in vivo* models, and in combination with mifamurtide, an immunomodulatory drug currently used in Europe to treat osteosarcoma patients, inhibits osteosarcoma metastasis in immune competent mice. This provides a novel strategy for activating the innate immune system to better combat metastatic progression.

## Results

### DDX3 is expressed in osteosarcoma patient samples and associated with worse overall survival

The clinical relevance of DDX3 has been established in EWS patients. Previous work established that DDX3 is expressed in various EWS and OS cell lines and PDX models^17^ and showed that high expression of DDX3 is correlated with worse event-free and overall survival in patients with EWS^26^. To establish the clinical relevance of DDX3 for osteosarcoma, we performed immunohistochemical analysis of a tissue microarray (TMA) consisting of OS patient samples, scoring DDX3 staining intensity from 0 (lowest expression) to +3 (highest expression) (Figure 1A). Among 77 patient samples, 69% exhibited high DDX3 expression (+2 or +3), while 18% demonstrated low DDX3 expression (+1). Only 12% of samples expressed no DDX3 (Figure 1B). Using annotated clinical data from these samples, we evaluated the association between DDX3 expression and overall survival. Patients with high DDX3 expression demonstrated a trend toward worse overall survival compared to those with low expression (p=0.14; Figure 1C). Although this did not reach statistical significance, likely due to the limited sample size (n=77), the observed trend is consistent with prior findings in EWS. These findings highlight the clinical significance of DDX3 and its potential as a therapeutic target to improve treatment outcomes for patients with bone sarcomas.

**Figure 1:**
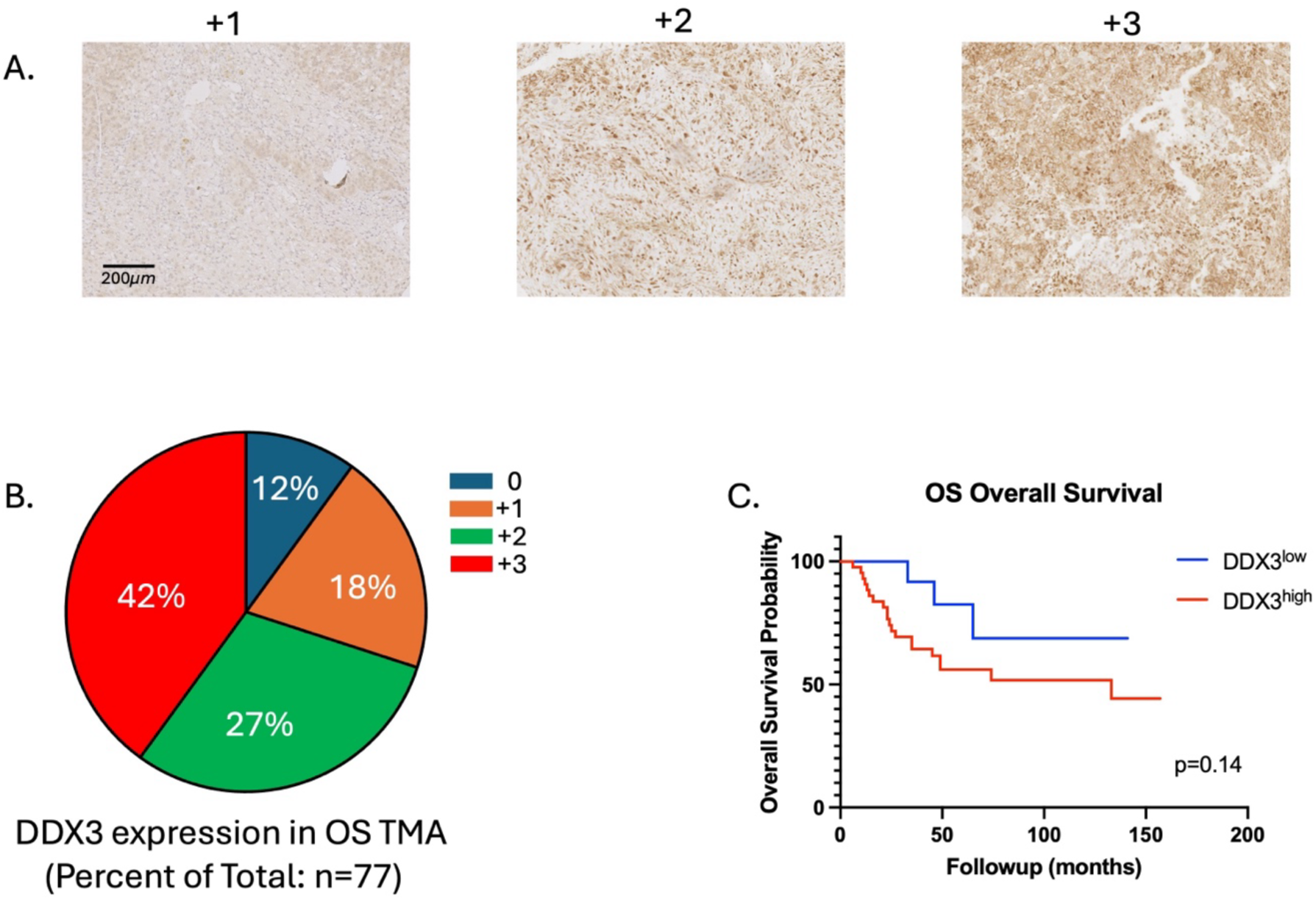
Elevated DDX3 expression in Osteosarcoma is associated with poor prognosis. (A) Immunohistochemical staining of osteosarcoma tissue microarray (TMA). Representative images of DDX3 protein expression. Mag bar =200*μm.* (B) Pie chart showing the distribution of DDX3 expression among TMA samples (n=77). (C) Kaplan-Meier curves demonstrating overall survival of osteosarcoma patients with either low (red) or high (blue) levels of DDX3 mRNA expression. p=0.14 by log rank (Mantel Cox) test.

### DDX3 inhibition leads to an accumulation of cytoplasmic double stranded RNAs

Previous work in breast cancer showed that DDX3 inhibition leads to an accumulation of cytoplasmic dsRNA which then triggers a downstream pathway culminating in a Type I interferon response^23^. To confirm that this is a common mechanism for controlling cytoplasmic nuclei acids in tumor cells, we quantified endogenous dsRNAs in both EWS and OS using the J2 monoclonal anti-dsRNA antibody, which has been previously validated and is widely used to recognize viral and cellular dsRNA^23, 27^. Three Ewing sarcoma cell lines (TC71, TC32, and A4573) (Figure 2A) and three osteosarcoma cell lines (HOS, SaOS2, and U2OS) (Figure 2C) were treated with RK-33 for 24 hours and evaluated by immunofluorescence using the J2 antibody. Inhibition of DDX3 by RK-33 caused a statistically significant accumulation of dsRNAs (Figure 2B and 2D). To confirm that the observed effect was specific to DDX3 inhibition and not due to off-target effects, we also treated DDX3 knockdown cell lines (M1F6 and M1F10^17^) alongside a scramble control (MSD10^17^) and the parent cell line TC71 (Figure 2E and Supplementary Figure 1A). RK-33 had no effect on cytoplasmic dsRNA in the DDX3 knockdown cell lines, supporting the specificity of RK-33 for DDX3 (Figure 2F). The knockdown cell lines showed increased cytoplasmic dsRNA compared with the scramble control (Figure 2G), consistent with DDX3 being the key modulator of dsRNA accumulation in the cytoplasm of sarcoma cells. In summary, in both Ewing sarcoma and osteosarcoma cell lines, inhibition of DDX3 activity by knockdown or RK-33 leads to an accumulation of dsRNAs, suggesting DDX3 is the primary regulator of endogenous cytoplasmic dsRNAs.

**Figure 2:**
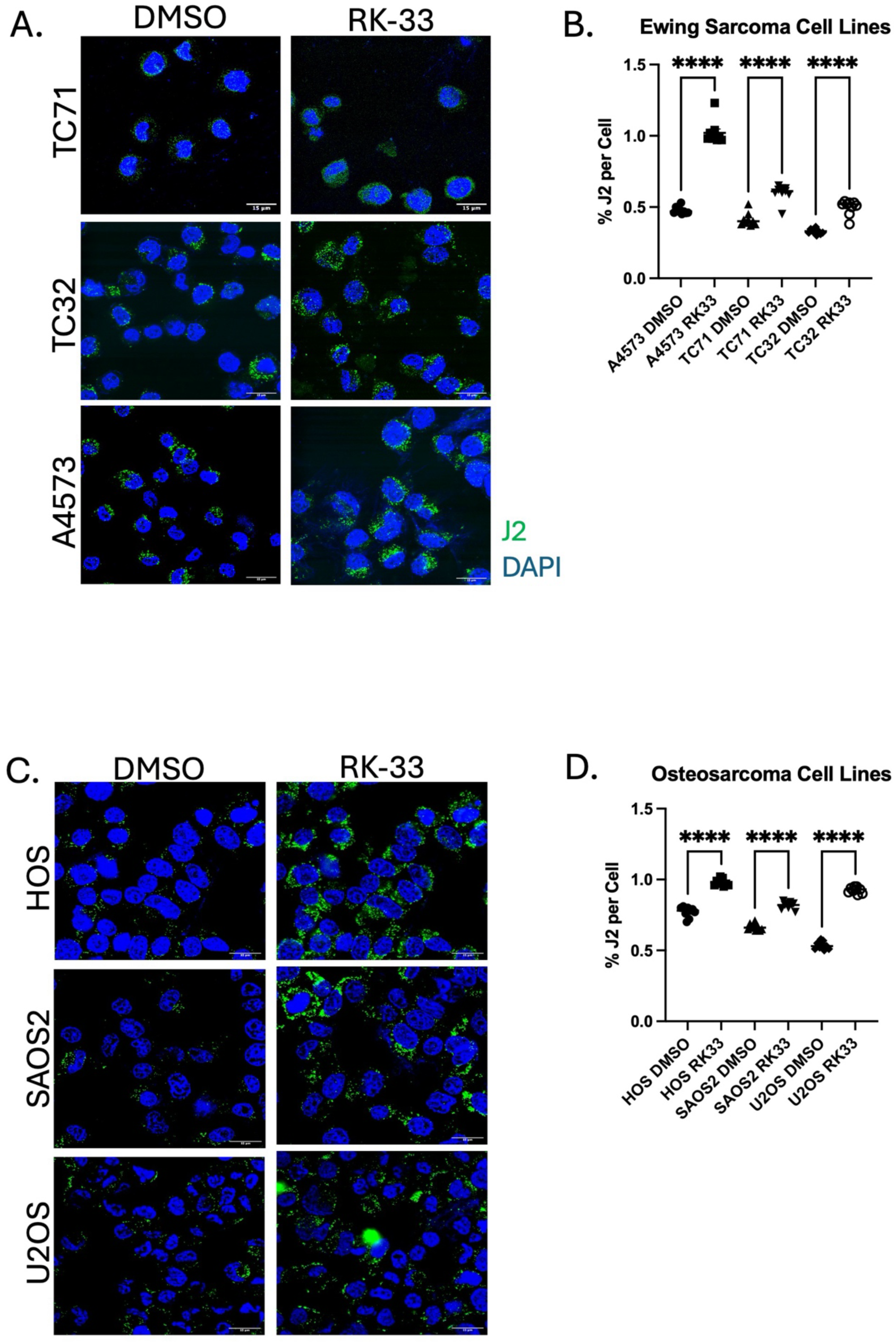
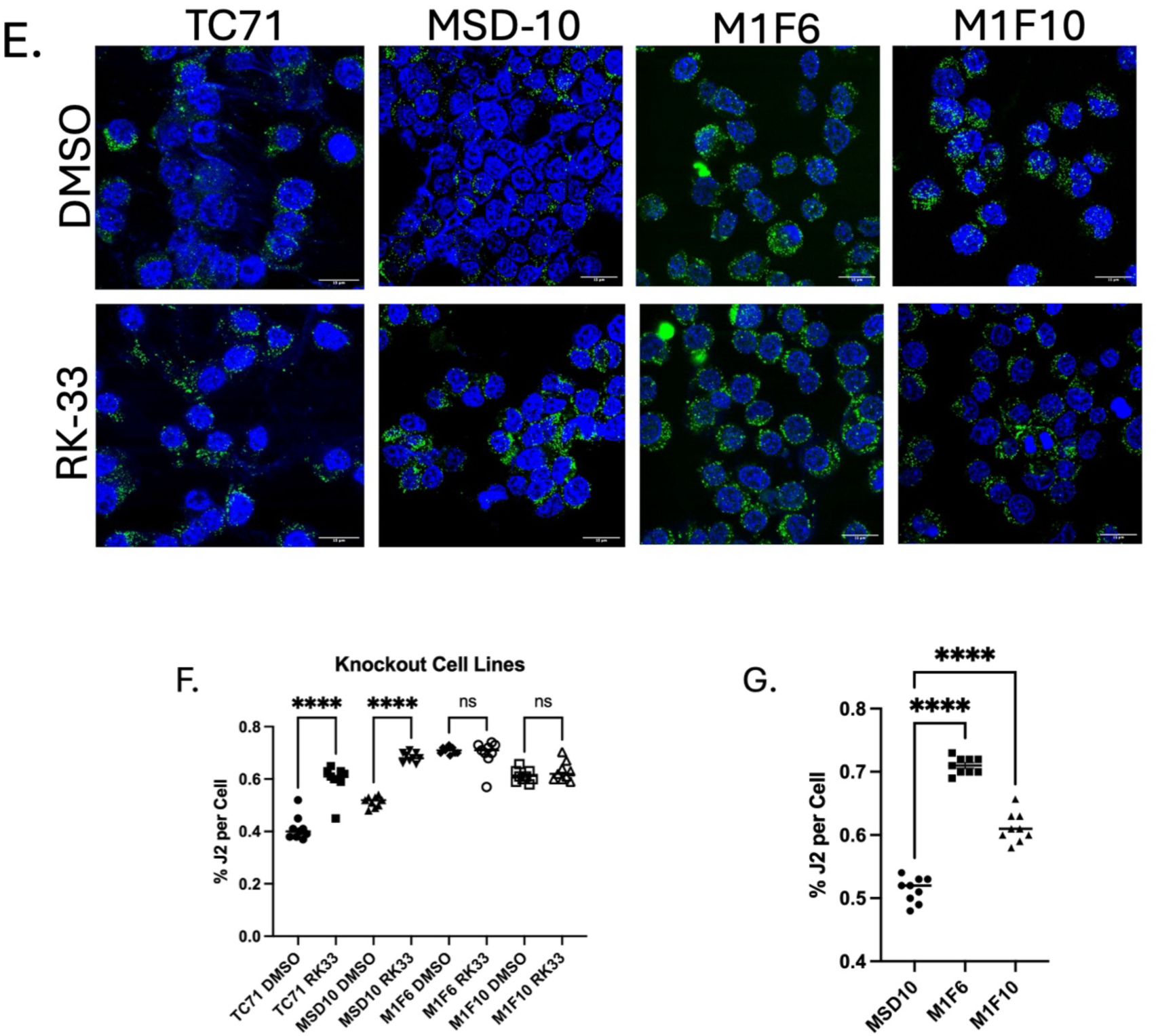
DDX3 inhibition leads to an accumulation of dsRNAs. (A) Ewing sarcoma TC71, TC32, and A4573 cell lines analyzed by immunofluorescence following 24-hour treatment with either RK-33 or DMSO for accumulation of dsRNA with anti-dsRNA specific antibody (J2). Mag=15*μm*. (B) Quantitative analysis of dsRNA accumulation. (C) Osteosarcoma HOS, SaOS2, and U2OS cell lines analyzed by immunofluorescence following 24-hour treatment with either RK-33 or DMSO for accumulation of dsRNA with anti-dsRNA specific antibody (J2). Mag=15*μm*. (D) Quantitative analysis of dsRNA accumulation. (E) DDX3 knockdown cell lines M1F6 and M1F10, scramble control MSD-10, and representative Ewing sarcoma cell line TC71 analyzed by immunofluorescence following 24-hour treatment with either RK-33 or DMSO for accumulation of dsRNA with anti-dsRNA specific antibody (J2). Mag=15*μm*. (F) Quantitative analysis of dsRNA accumulation. (G) Quantitative analysis of dsRNA accumulation in knockdown cell lines at baseline exposure, prior to any treatment. Error bars are the SEM of triplicate experiments, and each experiment was repeated three times. ****=p<0.0001.

### DDX3 inhibition induces a Type I interferon and innate immune response

To determine if the accumulation of cytoplasmic dsRNA induces a Type I interferon response in bone sarcoma cells, we evaluated the expression of a panel of genes divided into three categories. Interferon stimulated genes (ISG): *IF144L, OAS1, OAS2, IFIT2, IFIT3, ISIG15, STAT1*, *IRF3, IRF5, IRF7*; antigen presentation pathway genes (APP): *PSMB8, TAP1, HLA-A, HLA-B, HLA-C, HLA-DRA*; and a group of cytokine/chemokine genes: *CCL5, IL6, IL1-B, CCL4, CXCL10*. Using qRT-PCR analysis we found an overall increased expression of the ISGs and cytokines/chemokines in response to RK-33 treatment in EWS and OS cell lines (Figure 3A and Supplementary Figure 2A). APP genes showed little change in response to RK-33 treatment, highlighting the specificity of which downstream pathways are regulated by DDX3.

**Figure 3:**
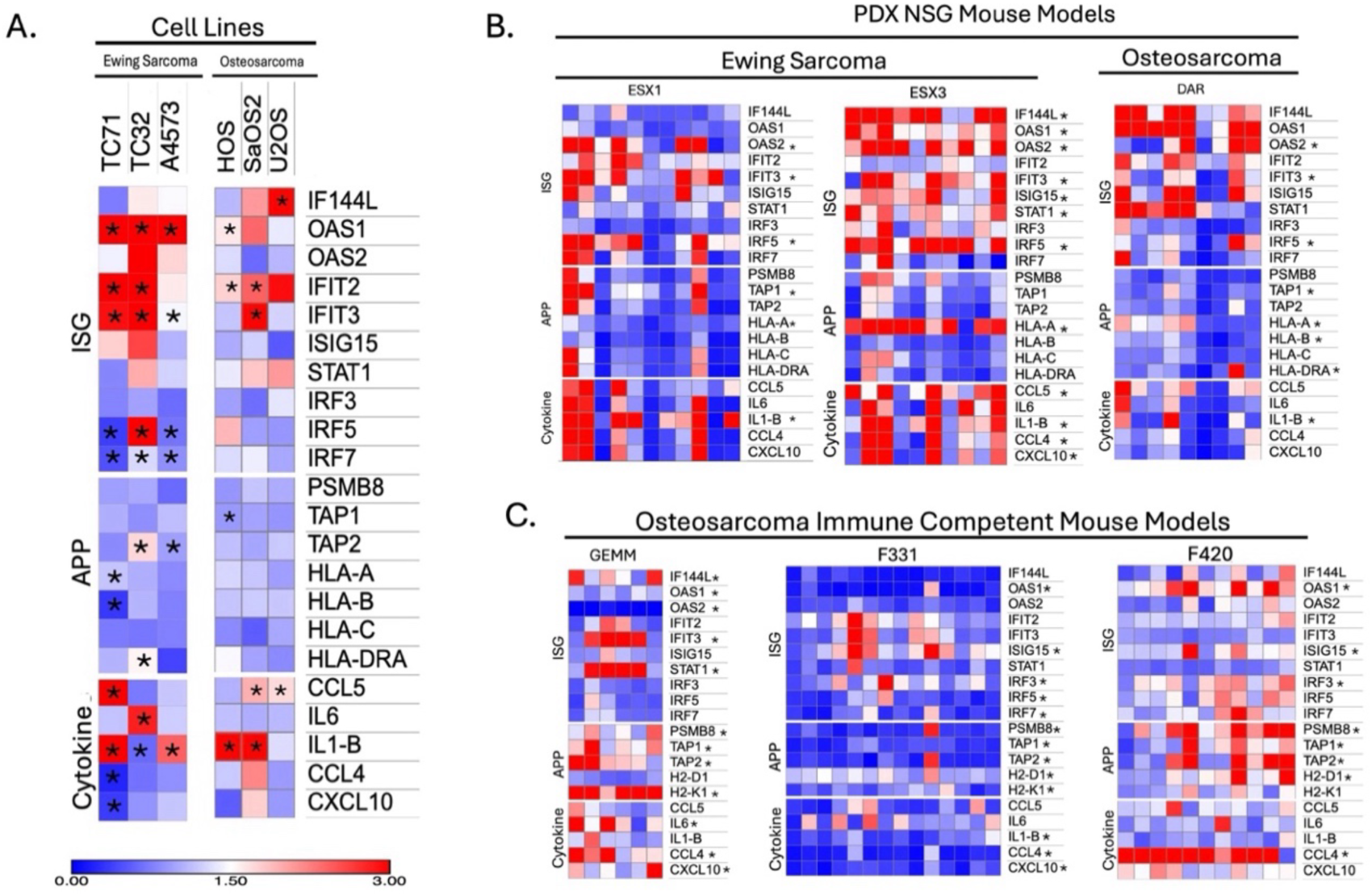
RK33 triggers interferon stimulating genes *in vivo and in vitro*. (A)The indicated cell lines were treated with RK-33 for 24 hours, mRNA was isolated, reverse transcribed, and anlyzed via qRT-PCR. Expression of the indicated gene in treated cells was quantified relative to control cells by the ΔΔCt method after normalization based on beta-2-microglobulin expression. A heatmap of the results is presented. The deepest blue represents downregulation in fold change expression levels in treated cells compared with control, and the deepest red represents upregulation in fold change expression levels in treated cells compared with control. Data is representative of triplicate experiments repeated independently three times, * indicates statistically significance (p < 0.05). (B) Using two different Ewing sarcoma patient derived xenografts (ESX1 and ESX3) as well as one osteosarcoma patient derived xenograft (DAR) implanted into NSG mice and treated with RK-33. Tumors were extracted and dissociated, mRNA was isolated, reverse transcribed, and analyzed via qRT-PCR. Data was analyzed as described in (A). A heatmap of the results is presented with each column representing an individual mouse (ESX1: n=11, ESX3: n=10, DAR: n=9). Data is representative of triplicate experiments, * indicates statistical significance (p < 0.05). (C) mRNA was isolated from tumors from three immune competent models of osteosarcoma (genetically engineered model of spontaneous OS, two syngeneic murine models F331 and MF420) treated with RK-33. Tumors were processed and mRNA was isolated, reverse transcribed, and analyzed via qRT-PCR as described in (A) and (B). A heatmap of the results is presented with each column representing an individual mouse (GEMM: n=6, F331: n=14, MF420: n=11). Data is representative of triplicate experiments, * indicates statistical significance (p < 0.05).

In addition to these *in vitro* studies, we evaluated the effect of RK-33 *in vivo* on mouse models of EWS and OS, using both immune deficient and immune competent models. Currently, there is no immune competent mouse model of Ewing sarcoma^28,29,30,31^, necessitating the use of immune deficient mice. We implanted fragments of a low passage, EWS PDX (JHHESX1 and JHHESX3) as well as an OS PDX (DAR) orthotopically into NSG mice and determined the effect of RK-33 on the expression of the same panel of genes. Treated tumors had generally increased expression of the ISGs and cytokines/chemokines, similar to what we saw in our *in vitro* studies, but with some upregulation of some of the APP genes including TAP1, HLA-A, and HLA-DRA (Figure 3B and Supplementary Figure 2B). We also investigated gene expression changes in three different immune competent models of osteosarcoma, including a genetically engineered mouse model (GEMM) in which loss of p53 and Rb causes spontaneous OS^32^ and two syngeneic murine osteosarcoma cell lines, F331 and F420 (Figure 3C), which were also derived from a genetically engineered model^28^. RK-33 induced gene expression changes in the GEMM and F420 models similar to the other models, with even more upregulation amongst the APP genes, likely because these are immune competent models (Supplementary Figure 2C). Interestingly, treatment with RK-33 induced significant downregulation across the entire gene panel in the F331 tumors.

While the targeted qPCR panel confirmed that DDX3 inhibition induces a Type I interferon response across multiple sarcoma models, it was limited to genes previously identified in breast cancer^23^ and could not capture the full transcriptomic impact of RK-33 in the sarcoma context, particularly evident by the unexpected downregulation observed in F331. To obtain a comprehensive view of how DDX3 inhibition modulates gene expression *in vivo,* we performed RNA sequencing across multiple sarcoma models. We implanted fragments of a low passage, EWS PDX (JHHESX1) orthotopically into NSG mice. For osteosarcoma, we used three different immune competent models, including a genetically engineered mouse model (GEMM) in which loss of p53 and Rb causes spontaneous OS^33^ and two syngeneic murine osteosarcoma cell lines, F331 and F420, which were also derived from a genetically engineered model^34^. Volcano plots identified genes differentially expressed between the control ESX1 tumors and those treated with RK-33 (Figure 4A), including *SERPINE1, CCL2, IFITM1, IER3, HLA-B, JUN, CXCL8,* and *TRIM8* (p < 0.05; Supplementary Table 3), and all play a role in stress response and innate immunity. Gene set enrichment analysis of ESX1 tumors treated with RK-33 (Supplementary Table 4) showed increased expression of genes associated with TNF-α signaling via NF-kB, interferon alpha response, interferon gamma response, and inflammatory response (Figure 4B). Volcano plots of genes differentially expressed between control GEMM tumors and those treated with RK-33 (Figure 4C) demonstrate that *Slit2, Nrp2, Spp1, Cav1, Met, Cxcl13,* and *Emilin2* are some of the most differentially expressed genes (p < 0.05; Supplementary Table 5), all of which are involved in pro-inflammatory signaling. RK-33-treated GEMM tumors show significant enrichment (Supplementary Table 6) in pathways including wound healing, response to cytokine, regulation of innate immune response, and Type I interferon mediated response (Figure 4D). In F331 tumors, differentially expressed genes following RK-33 treatment included *Enpp2, Fes,* and *Ndrg2* (p<0.05; Supplementary Table 7), genes associated with inflammatory regulation and macrophage recruitment (Figure 4E). Pathway enrichment revealed upregulation of IL2 STAT5 signaling, inflammatory pathways, and TNF-α signaling via NF-kB (Figure 4F, Supplementary Table 8). Notably interferon response pathways were downregulated in F331, consistent with that targeted qPCR panel. In F420 tumors, volcano plot analysis identified enrichment of genes including *Cxcl5, Serpine, Ccr5,* and *CCL12* (p<0.05; Supplementary Table 9), all involved in immune cell recruitment (Figure 4G), with pathway enrichment analysis identifying upregulation of interferon gamma and alpha response and inflammatory response pathways (Figure 4H; Supplemetary Table 10). Taken together, RNA sequencing data demonstrated that DDX3 inhibition via RK-33 consistently induces pro-inflammatory and innate immune activation signatures across multiple bone sarcoma models *in vivo*.

**Figure 4:**
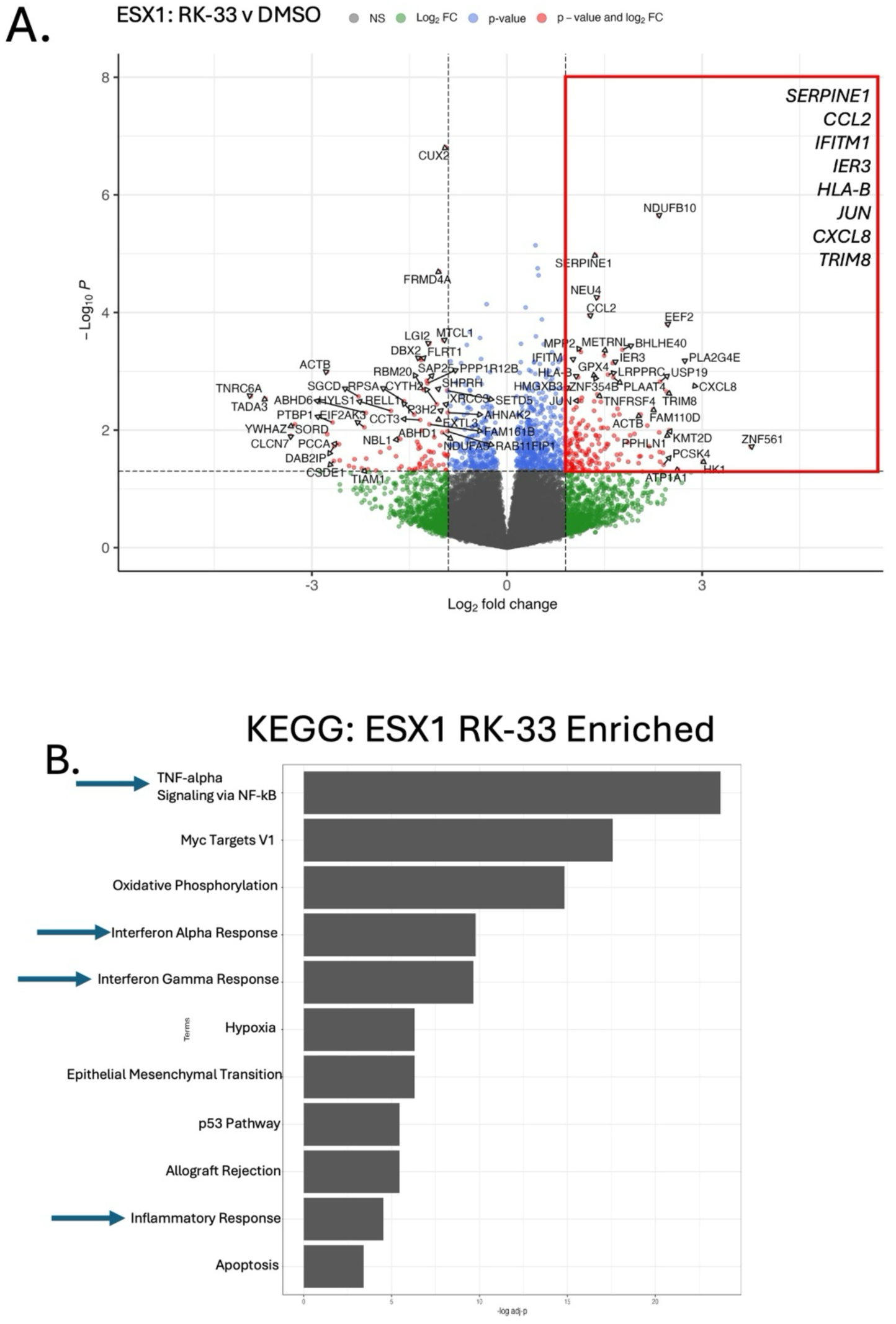
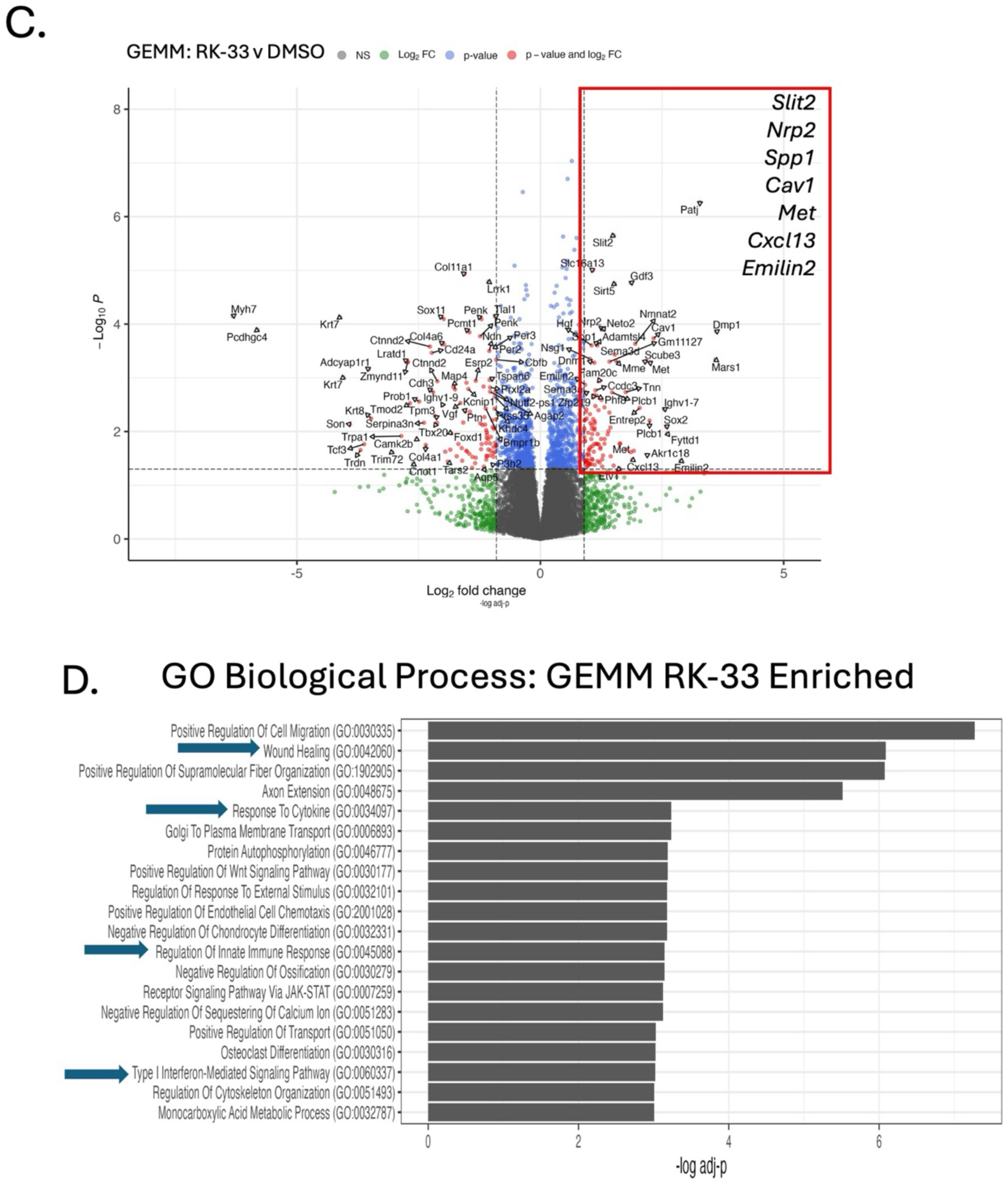
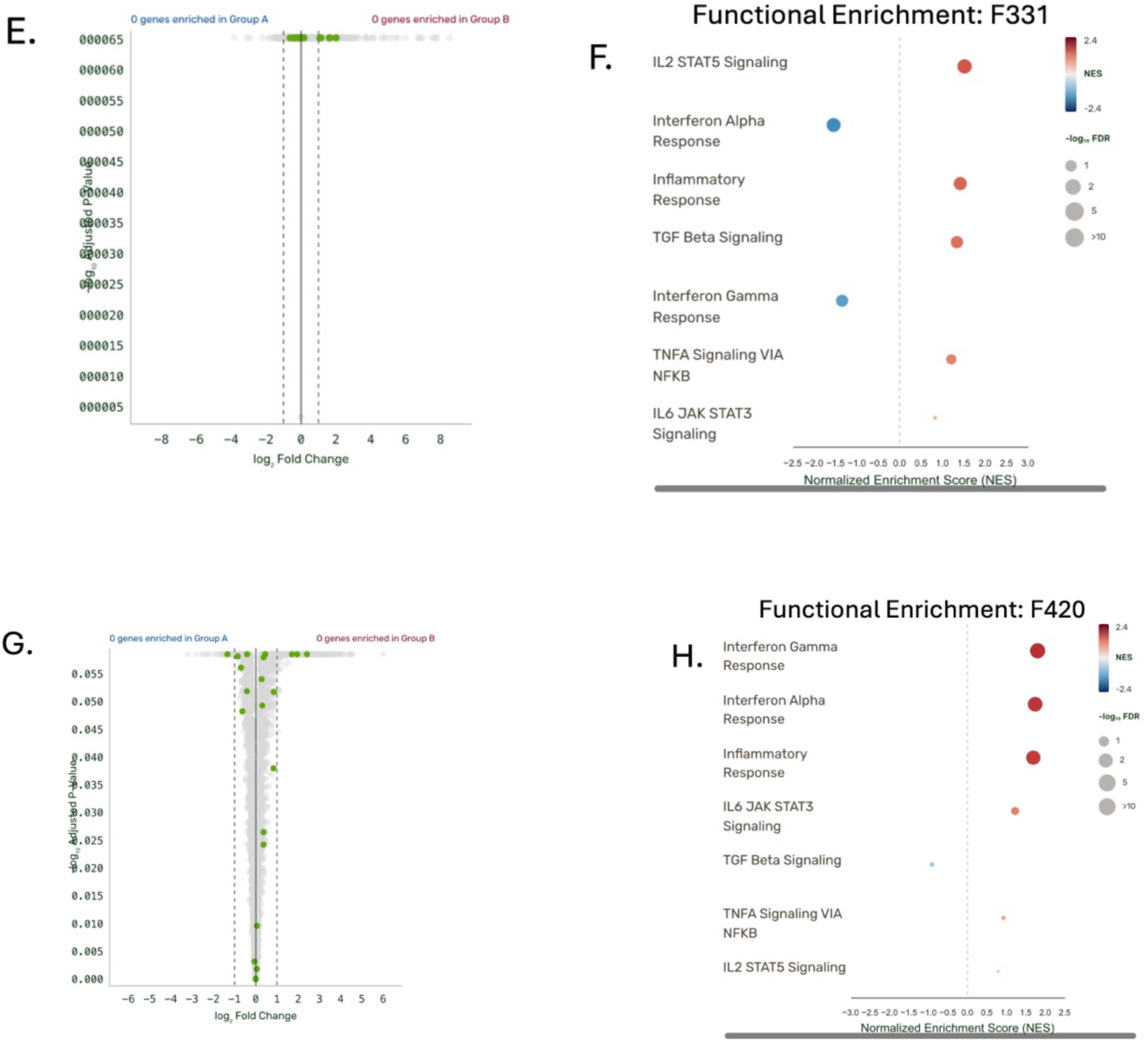
Transcriptomic analysis demonstrates RK33 triggers immune related pathways in vivo. (A) Volcano plot depicting differentially expressed genes in ESX1 tumor cells. Upper right quadrant (red box) represents genes upregulated in response to RK-33 treatment (p-value < 0.05). Left upper quadrant represents genes downregulated in response to RK-33 treatment (p-value < 0.05). Relevant upregulated genes in response to RK-33 include *SERPINE1, CCL2, IFITM1, IER3, HLA-B, JUN, CXCL8, TRIM8.* (B) KEGG analysis showed upregulation of genes involved in TNF-alpha signaling via NF-kB, interferon alpha response, interferon gamma response, and inflammatory response. (C) Volcano plot depicting differentially expressed genes in GEMM tumor cells. Upper right quadrant (red box) represents genes upregulated in response to RK-33 treatment (p-value < 0.05). Left upper quadrant represents genes downregulated in response to RK-33 treatment. Relevant upregulated genes in response to RK-33 include *Slit2, Nrp2, Spp1, Cav1, Met, Cxcl13, Emilin2.* (D) Enriched biological processes and upregulation of genes involved in wound healing, response to cytokine, regulation of innate immune response, and type I interferon-mediated signaling pathway. (E)Volcano plot depicting differentially expressed genes in F331 tumor cells. (F) Pathway enrichment analysis from F331 tumors show upregulation of IL2 STAT5 signaling, inflammatory response, TGF beta signaling, and TNF-α signaling via NF-kB. (G) Volcano plot depicting differentially expressed genes in F420 tumor cells. (H) Pathway enrichment analysis from F420 tumors show upregulation of interferon gamma and alpha response, inflammatory response and IL6 JAK STAT3 signaling.

### RK-33 treatment induces an accumulation of pro-inflammatory type macrophages

We next aimed to confirm that the pro-inflammatory changes in the TME caused by DDX3 inhibition alters macrophage infiltration and polarization using immunofluorescence staining focused on established surface markers specific for M1 and M2 macrophage subtypes. RK-33 treatment significantly increased the number of Iba1^+^ macrophages expressing the M1 marker CD86^+^ and decreased the number of cells with the M2 marker Arg1^+^ in ESX3 tumors grown in NSG mice compared to the controls (Figure 5A and 5B). F331 tumors similarly responded to RK-33 with a significant increase in CD86^+^/Iba1^+^ staining and a reduction in Arg1^+^/Iba1^+^ cells (Figure 5C and 5D). In contrast, F420 tumors exhibited the opposite pattern, with a decreased number of M1 macrophages and an increased number of M2 macrophages in the RK-33 treated tumors (Figure 5E and 5F).

**Figure 5:**
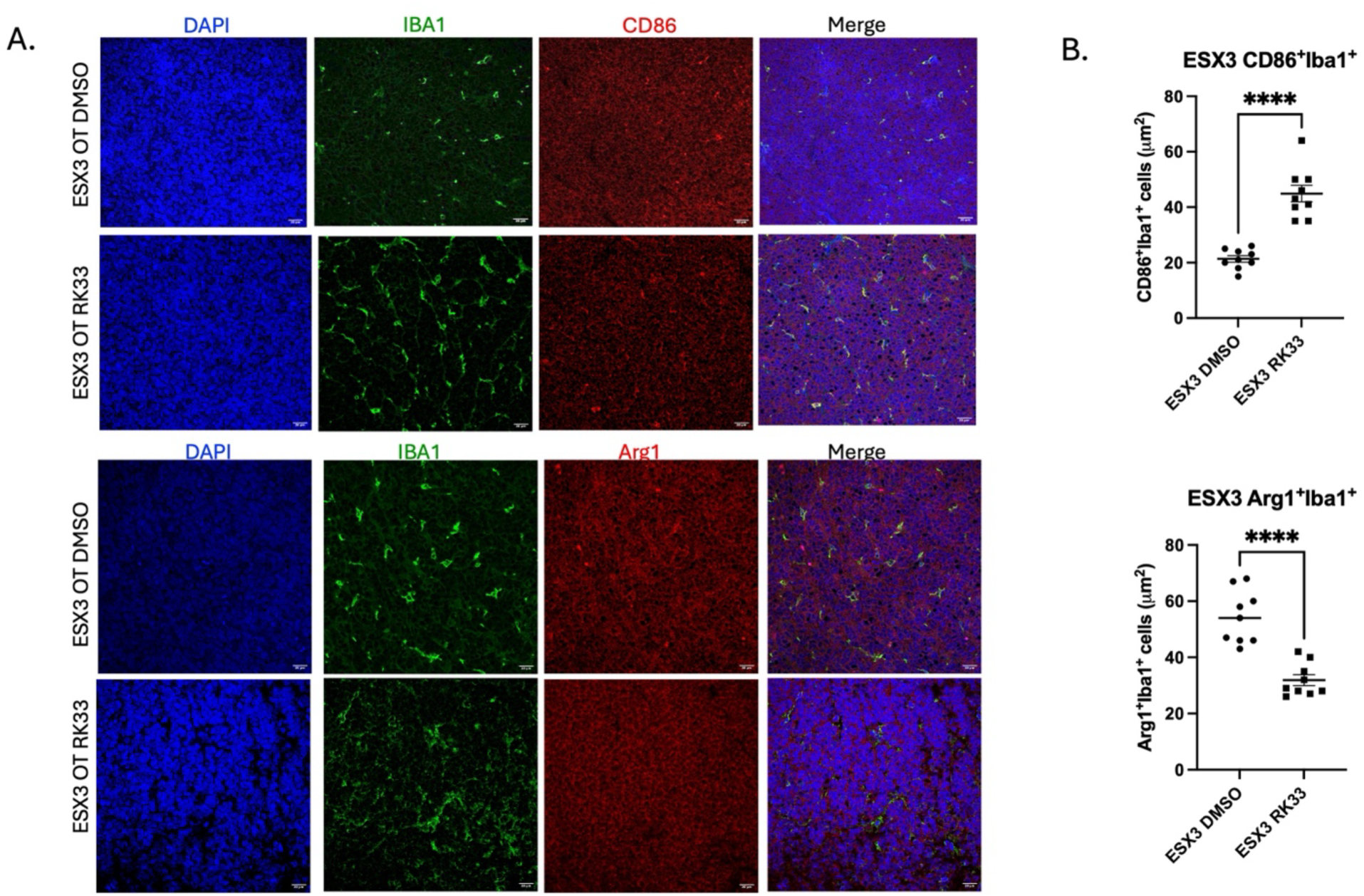
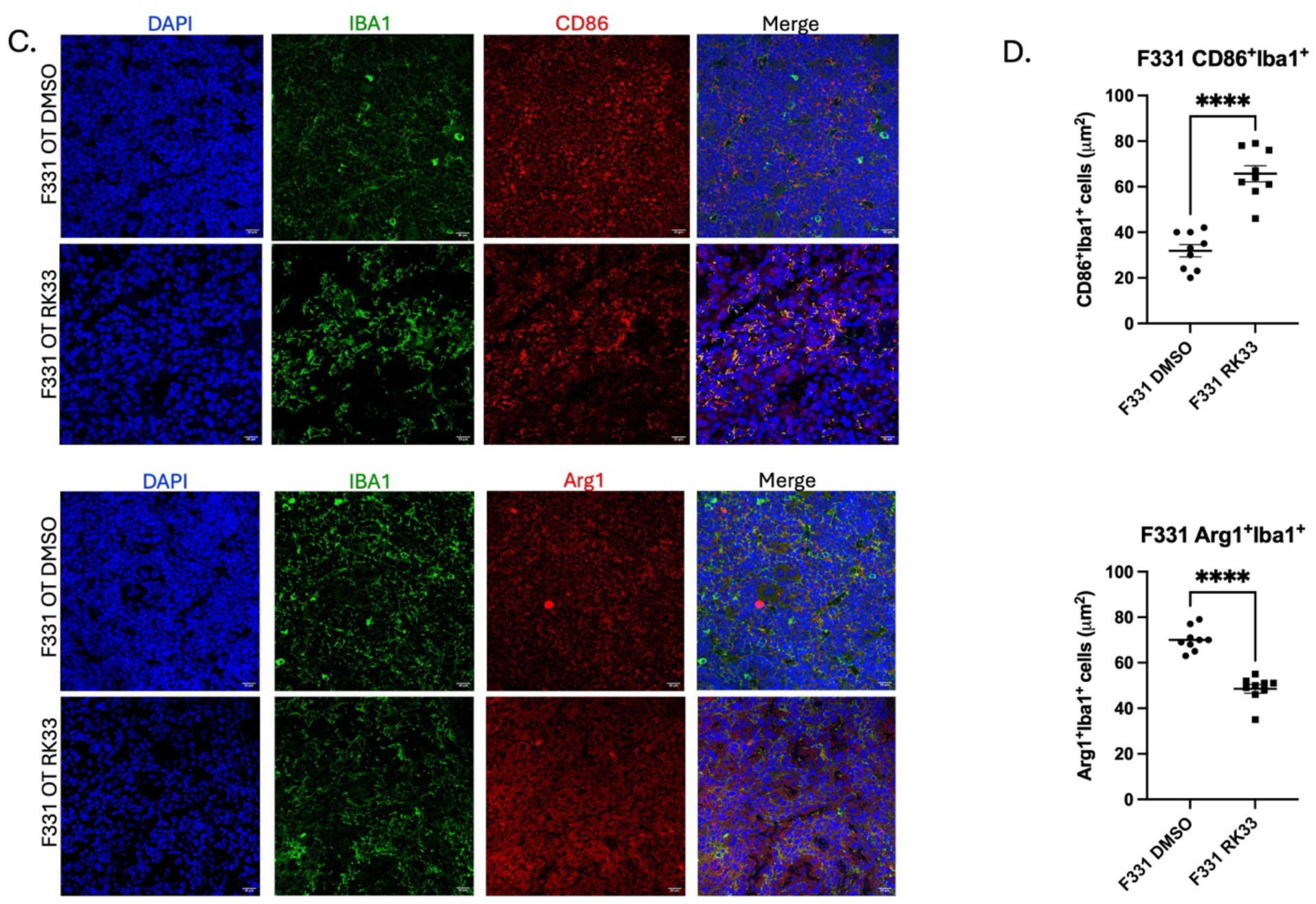
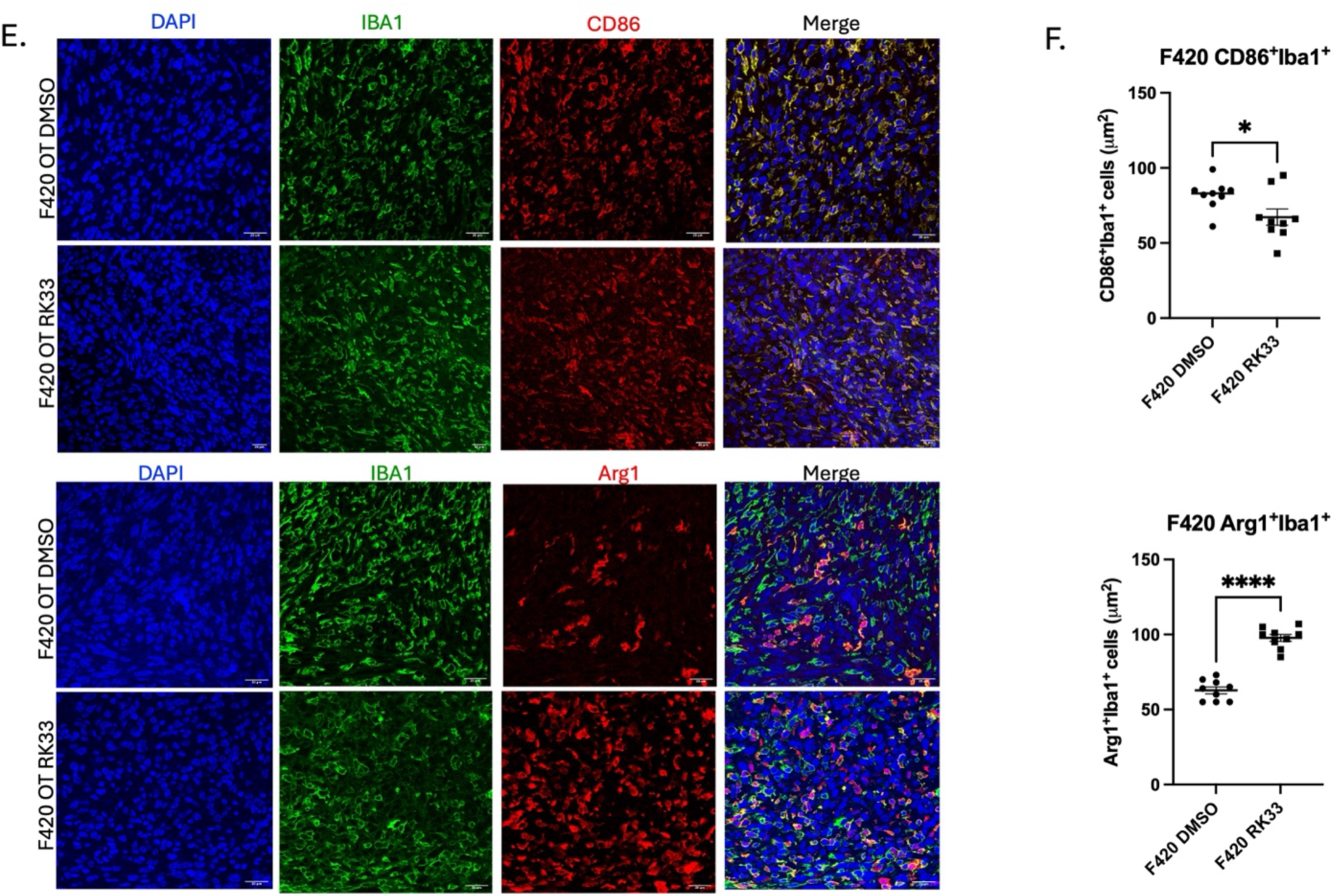
DDX3 inhibition leads to an accumulation of pro-inflammatory macrophages. (A)Representative photomicrographs of immunofluorescence analysis of macrophage polarization from ESX3 tumors treated with either RK-33 or DMSO. M1 phenotype: CD86+ (red) and Iba1+ (green); M2 phenotype: Arg1+ (red) and Iba1+ (green). Mag=20*μm*. (B) Quantitative analysis of M1 macrophage phenotype (CD86^+^Iba1^+^) (p<0.0001) and M2 macrophage phenotype (Arg1^+^Iba1^+^) (p<0.0001). (C) Representative photomicrographs of immunofluorescence analysis of macrophage polarization from F331 tumors treated with either RK-33 or DMSO. M1 phenotype: CD86+ (red) and Iba1+ (green); M2 phenotype: Arg1+ (red) and Iba1+ (green). Mag=20*μm*. (D) Quantitative analysis of M1 macrophage phenotype (CD86^+^Iba1^+^) (p<0.0001) and M2 macrophage phenotype (Arg1^+^Iba1^+^) (p<0.0001). (E) Representative photomicrographs of immunofluorescence analysis of macrophage polarization from F420 tumors treated with either RK-33 or DMSO. M1 phenotype: CD86+ (red) and Iba1+ (green); M2 phenotype: Arg1+ (red) and Iba1+ (green). Mag=20*μm*. (F) Quantitative analysis of M1 macrophage phenotype (CD86^+^Iba1^+^) (p=0.037) and M2 macrophage phenotype (Arg1^+^Iba1^+^) (p<0.0001). Error bars are the SEM of triplicate experiments, and each experiment was repeated three times.

Although immunofluorescence can provide a broad view of macrophage polarization in tumors, there is a limit to the number of markers that can be evaluated in a single experiment. To provide a more detailed view, we turned to cytometry by time of flight (CyTOF). This high-dimensional mass cytometry technique labels antibodies with purified heavy-metal tags that biological systems do not normally contain, allowing the simultaneous detection of 50 or more markers in a single sample^35^. This capability provides an advantage over flow cytometry, which relies on fluorophore-labeled antibodies to detect cellular targets and therefore limits the number of markers that can distinguished without excessive spectral overlap or autofluorescence^36^. Using our specially designed CyTOF panel, we evaluated ESX3 (EWS), F331(OS), and F420 (OS) tumors in mice treated with or without RK-33 to characterize infiltrating macrophage populations. Within the ESX3 tumors, high-dimensional UMAP visualization revealed distinct clusters of tumor associated macrophages based on cell surface marker expression. Breaking down the tumor population, the cells were divided into TAM negative cells (CD45^+^F4/80^−^, green) and TAM positive cells (CD45^+^F4/80^+^, cyan). Within the TAM positive population, specific subpopulations were categorized as M1-like (brown) or M2-like (purple) (Figure 6A). Treatment with RK-33 induces a noticeable shift in the density and distribution of these clusters, particularly within the M2 population, with a decrease in these markers in the RK-33 treated tumors (Figure 6B). Individual markers were quantified, and while there is a shift within the M1 population, the M2 population (M2 like/MHC II, LAP-TGF, CD204/SR, TLR1) showed a statistically significant decrease within the DDX3 inhibited tumors (Figure 6C). To ensure the presence of functional macrophages, ESX3 analysis was conducted in nude mice rather than NSG mice. The concordance between these results and those observed in our immunofluorescence experiments suggests that while neither model fully recapitulates an immune competent host, both support sufficient macrophage activity for meaningful polarization analysis. F331 tumor UMAP plots also reveal a discernible decrease of the M2-like cluster (cyan) and a corresponding expansion in the density of the M1-like population (dark green) in the RK-33 treated tumors compared to the control (Figure 6D). RK-33-treated tumors show consistent depletion of the M2 subtype across the polarization spectrum (Figure 6E).

**Figure 6:**
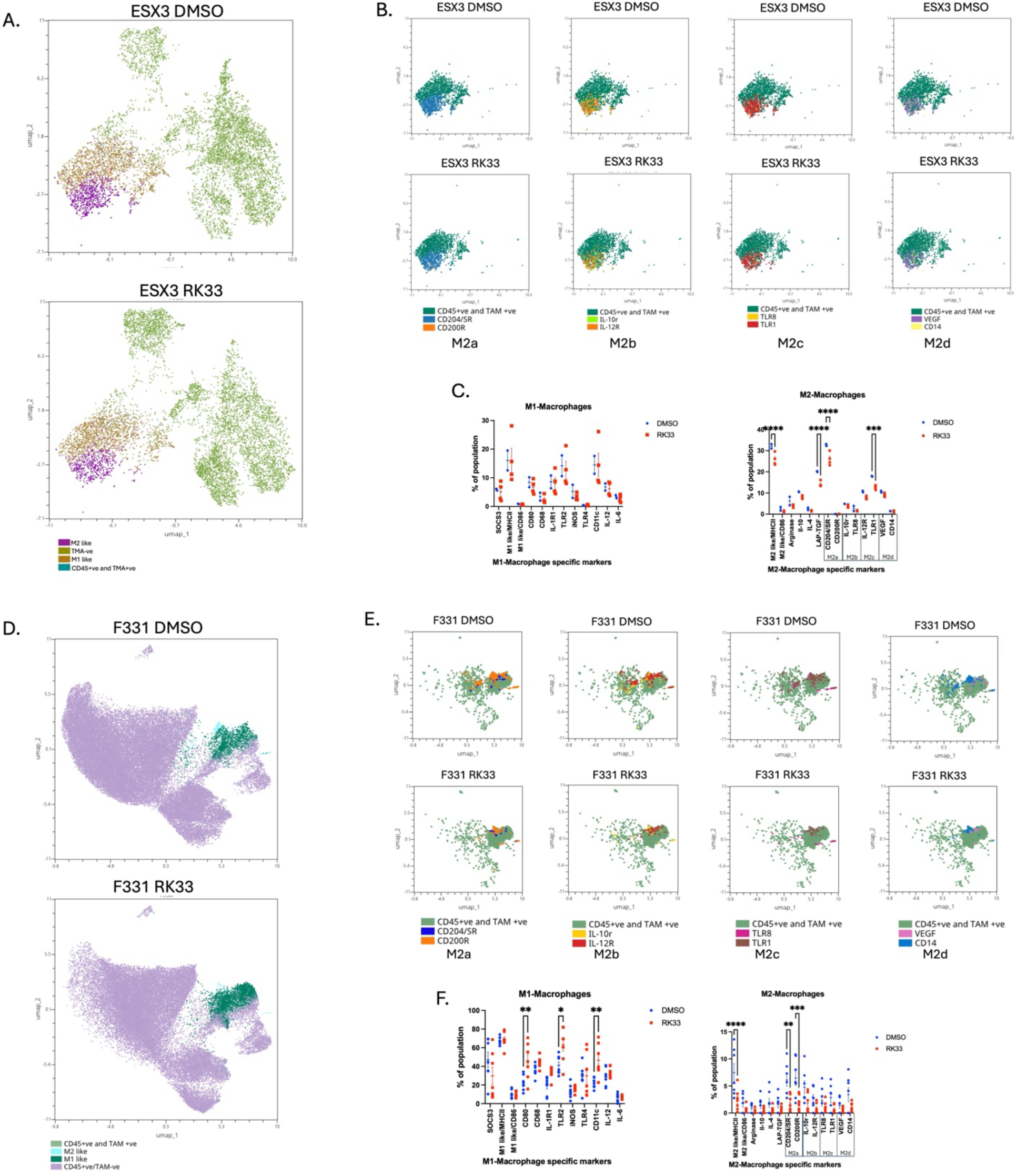
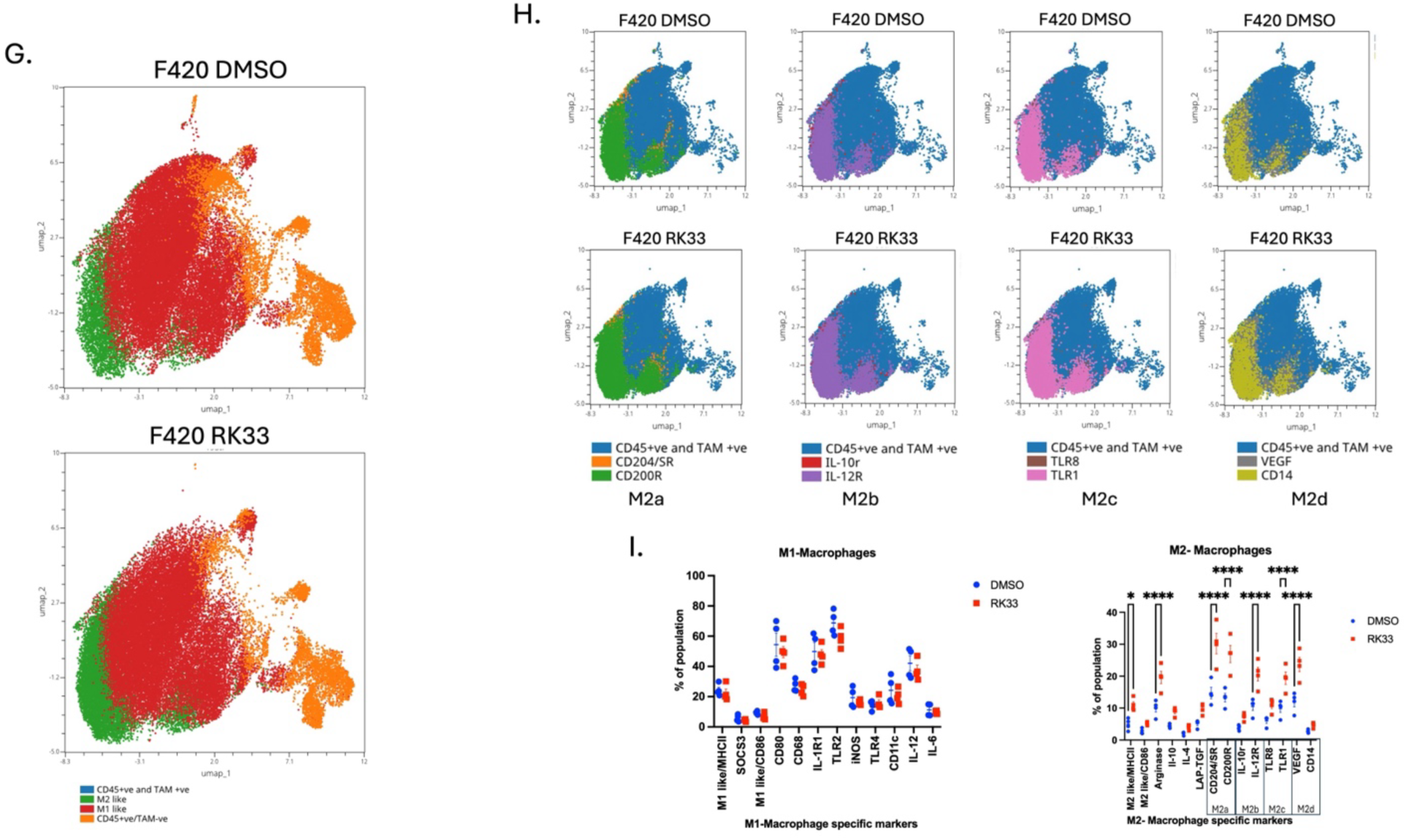
CyTOF analysis of macrophage polarization following RK33 treatment. (A) UMAP analysis of the TAM+ populations further classified into M1 like and M2 like populations in ESX3 tumors treated with RK-33 or DMSO. (B) UMAP plots showing the distribution of different macrophage markers representing the various stages of M2 like macrophages in the CD45+TAM+ populations in ESX3 tumors treated with RK-33 or DMSO. (C) Quantification of the breakdown in macrophage type population between the treated and control tumors. (M2like/MHCII p<0.0001; LAP-TGF p<0.0001; CD204/SR p<0.0001; TLR1 p=0.0001). (D) UMAP analysis of the TAM+ populations further classified into M1 like and M2 like populations in primary F331 tumors treated with RK-33 or DMSO. (E) UMAP plots showing the distribution of different macrophage markers representing the various stages of M2 like macrophages in the CD45+TAM+ populations in F331 tumors treated with RK-33 or DMSO. (F) Quantification of the breakdown in macrophage type population between the treated and control tumors. (M1: CD80 p=0.0053; TLR2 p=0.0105; CD11c p=0.003) (M2: M2like/MHCII p<0.0001; CD204/SR p=0.0011; CD200R p=0.0003). (G) UMAP analysis of the TAM+ populations further classified into M1 like and M2 like populations in primary F331 tumors treated with RK-33 or DMSO. (H) UMAP plots showing the distribution of different macrophage markers representing the various stages of M2 like macrophages in the CD45+TAM+ populations in F331 tumors treated with RK-33 or DMSO. (I) Quantification of the breakdown in macrophage type population between the treated and control tumors. (M2like/MHCII p=0.0320; Arginase p<0.0001; CD204/SR p<0.0001; CD200R p<0.0001; IL-12R p<0.0001; TLR1 p<0.0001; VEGF p<0.0001).

Quantitative analysis confirms a statistically significant increase in the M1 population (CD80, TLR2, CD11c) and a simultaneous statistically significant decrease in M2 populations (M2 like/MHC II, CD204/SR, CD200R) in treated tumors (Figure 6F). Interestingly, UMAP plots for F420 tumors identify a denser M2-like population (green) and a sparser M1-like cluster (red) in the RK-33 treated tumors (Figure 6G), consistent with our immunofluorescence results. Notably, RK-33 treatment was associated with a significant increase in the percentage of cells expressing M2 subtype markers (M2 like/MHC II, Arginase, CD200R, IL-12R, TLR1, VEGF) (Figure 6H and 6I). While the ESX3 and F331 tumors exhibit results similar to our RT-PCR and RNA-seq data, a decrease in pro-tumorigenic macrophages indicative of a more inflammatory TME, the F420 model exhibits an enrichment of M2 macrophages contributing to an immunosuppressive environment.

### MYC overexpression distinguishes F420 from F331

The opposing macrophage response to RK-33 observed in F331 and F420, despite both cell lines showing an upregulation in inflammatory related pathways, raises the question of what differences between these two cell lines drives such divergent outcomes. Because both cell lines are derived from the same genetically engineered osteosarcoma model, we initially hypothesized that this differential response could be attributed to differences in DDX3 expression. Western blot analysis confirmed comparable DDX3 protein levels in both cell lines (Supplementary Figure 3A), and both demonstrated increased dsRNA accumulation following RK-33 treatment (Supplementary Figure 3B). Having ruled out DDX3 expression as the source of discrepant response, we sought to identify the molecular basis for this observation through transcriptomic profiling. RNA sequencing revealed considerable transcriptomic differences between F331 and F420, with genes upregulated in F420 showing strong enrichment for canonical MYC target genes, many of which exhibited both high fold change and strong statistical significance (Figure 7A), including *MYC, MAX, NPM1, NOLC1, PES1, GNL3, MCM5, PCNA, HK2,* and *IMPDH2* (Supplementary Table 11). Consistent with this, gene set enrichment analysis demonstrated robust enrichment of MYC target gene sets in F420, with elevated expression across multiple *MYC-*associated genes (Figure 7B) (Supplementary Table 12). Given the prominence of *MYC*-associated transcriptional programs in F420, we assessed MYC expression directly. Quantitative RT-PCR analysis revealed a statistically significant increase in *MYC* mRNA levels in F420 relative to F331, and Western blot analysis demonstrated significantly higher MYC protein expression in F420 compared to F331 (Figure 7C, D, and E). These data identify MYC overexpression as a defining molecular feature that distinguishes F420 from F331 and provides a molecular basis for the divergent macrophage responses observed following RK-33 treatment. Given that the efficacy of macrophage-targeting immunotherapies is highly dependent on the polarization state of tumor associated macrophages, we next evaluated whether these divergent responses would translate into differential therapeutic outcomes.

**Figure 7:**
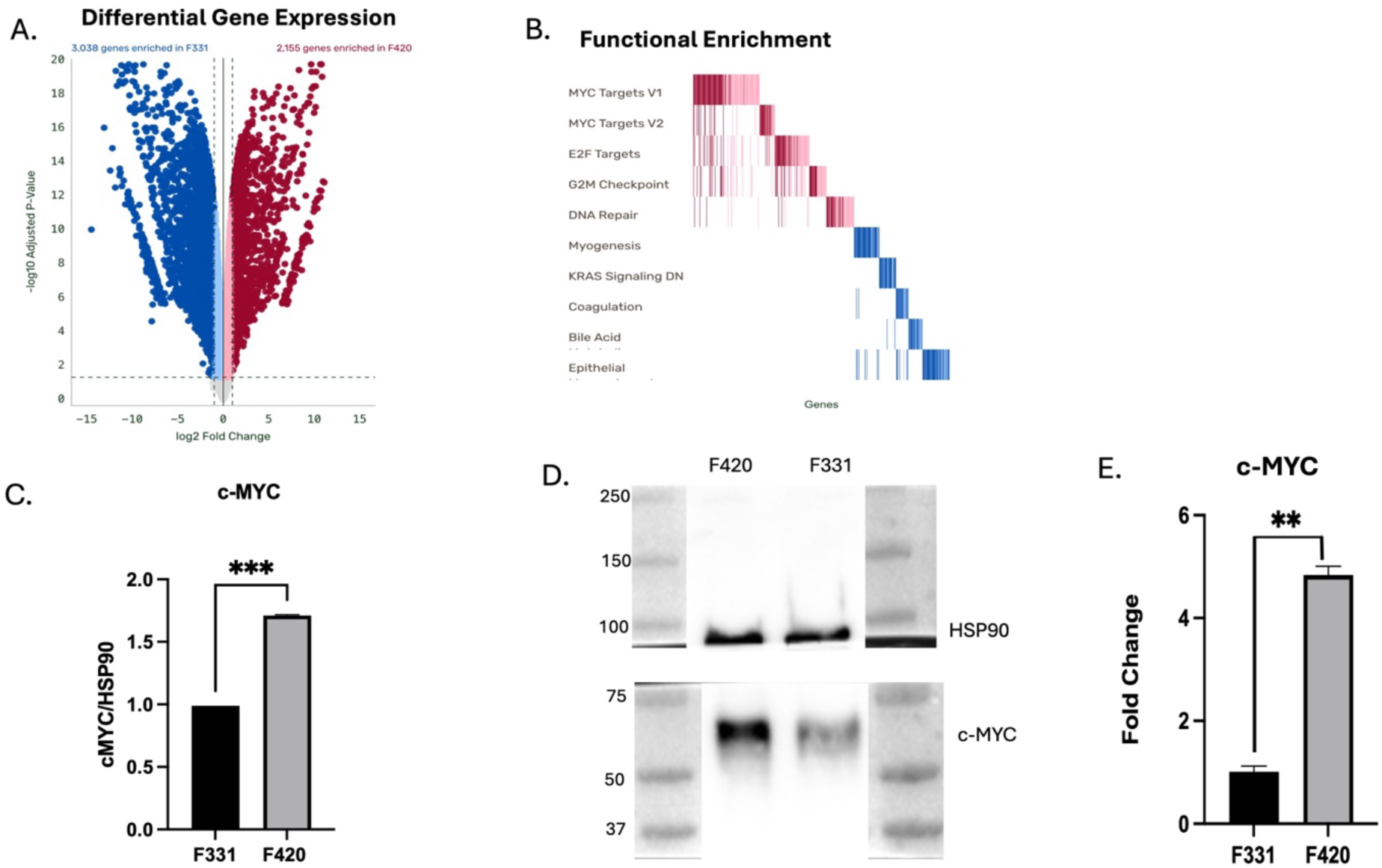
Comparing F331 and F420 cell lines. (A) Volcano plot depicting differentially expressed genes in the F331 and F420 cell lines (p-value < 0.05). (B) Hallmark gene sets derived from volcano plot confirming Myc targets as the top gene enrichment in the F420 cell line. (C) c-MYC mRNA expression quantified by qRT-PCR showed a statistically significant (p=0.0029) increased expression in the F420 cell line. (D) Western blot confirming higher expression of c-MYC in F420 cell line compared to F331. HSP90 was used as a loading control. (E) Quantification of western blot showing a statistically significant (p=0.0006) increase in protein expression in the F420 cell line compared to F331.

### Combination treatment with RK-33 and Mifamurtide inhibits OS metastases

Our data demonstrate that RK-33 modulates the response of the innate immune system to Ewing sarcoma and osteosarcoma tumors, with the induction of a Type I interferon response and a pro-inflammatory shift in intratumoral macrophage infiltration. Mifamurtide is an immunomodulatory drug, approved by the European Medicines Agency (EMA) for the treatment of osteosarcoma, that works by inducing expression of pro-inflammatory cytokines resulting in activation of tissue resident macrophages^37^. The ability of RK-33 to induce a pro-inflammatory macrophage polarization and the ability of mifamurtide to activate tissue resident macrophages suggest the possibility that these agents might be more effective together than individually. We utilized F331 and F420 tumors in our mouse model of spontaneous distant metastasis^38^. Tumor fragments were implanted into the tibias of C57BL/6 mice (Figure 8A). Upon confirmation of tumor engraftment, the mice were treated with either RK-33, mifamurtide, RK-33+mifamurtide, or vehicle control. When tumors reached a maximal diameter of 15 mm, the tumor-bearing limb was amputated, and treatment continued for eight more weeks, at which point the mice were euthanized and assessed for metastatic lesions upon necropsy. The F331 tumors, which respond to RK-33 with an inflammatory macrophage polarization, showed a treatment-dependent decrease in metastasis, with a decreased metastatic burden in mice treated with either RK-33 or mifamurtide, and a significantly decreased number of metastases in mice treated with the combination (p=0.03) (Figure 8B). Mice in the combination treatment cohort exhibited a statistically significant increase in proinflammatory CD86^+^/Iba1^+^ (Figure 8C and 8D) and decrease in anti-inflammatory Arg1^+^/Iba1^+^ (Figure 8E and 8F) macrophages compared to the controls. In contrast, mice implanted with F420 tumors, which have the opposite response to DDX3 inhibition, showed no difference in metastatic burden between the control and dual treated mice (p=0.9) (Figure 8G), and analysis of macrophage infiltration revealed a decrease in M1 macrophages (Figure 8H and 8I) and an increase in M2 macrophages (Figure 8J and 8K) in RK-33+mifamurtide treated tumors compared to controls. Thus, combination therapy improved outcomes in an OS model characterized by increased inflammatory macrophage infiltration in response to DDX3 inhibition, but not in a model that responds with increased anti-inflammatory macrophages. Together, these findings indicate that responses of these models to DDX3 inhibition differ markedly. Specifically, therapeutic efficacy in the F331 model is associated with the induction of a pro-inflammatory macrophage phenotype, whereas F420 tumors fail to mount a similar response and instead exhibit a shift toward an anti-inflammatory macrophage state following treatment.

**Figure 8:**
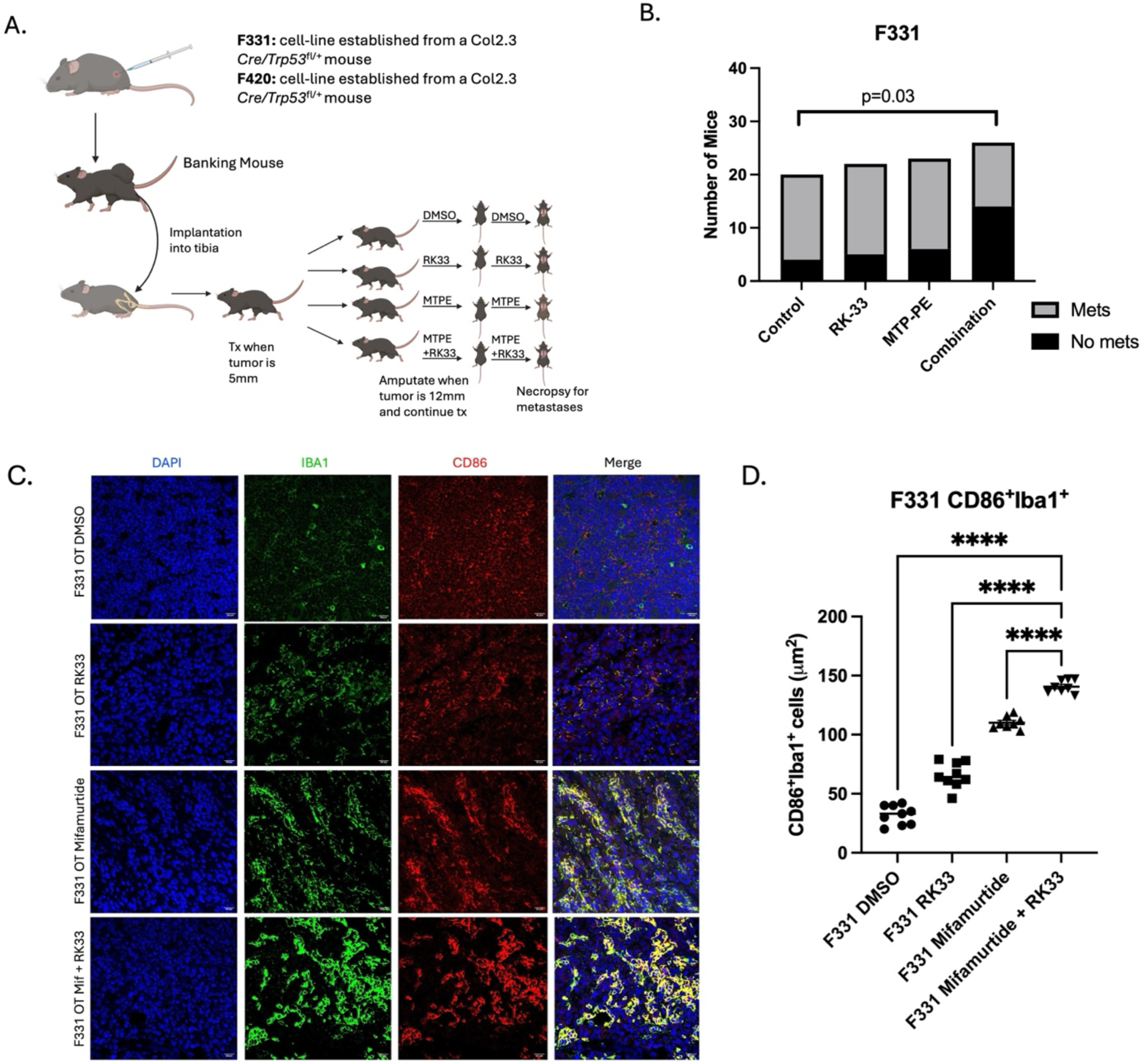
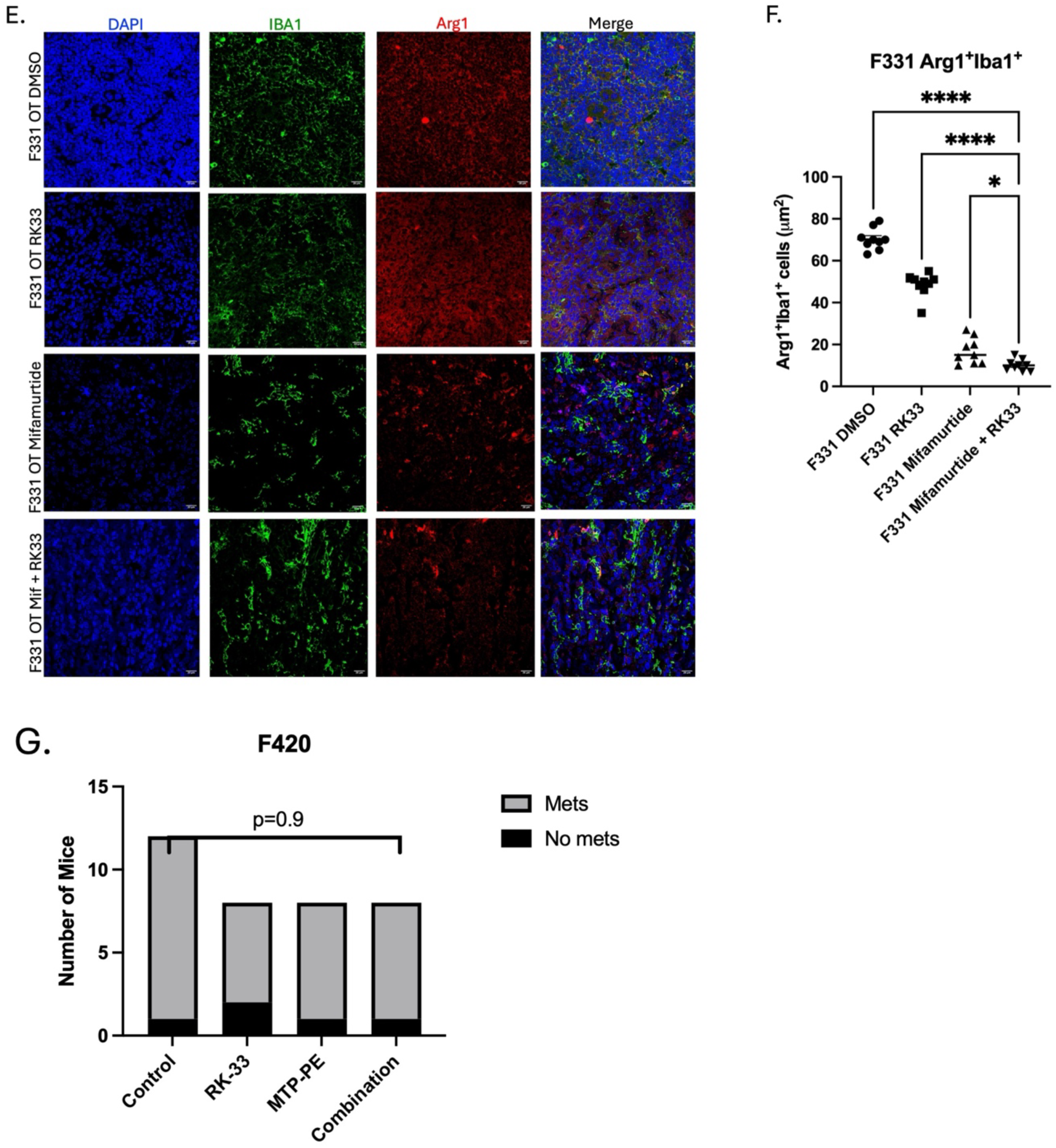
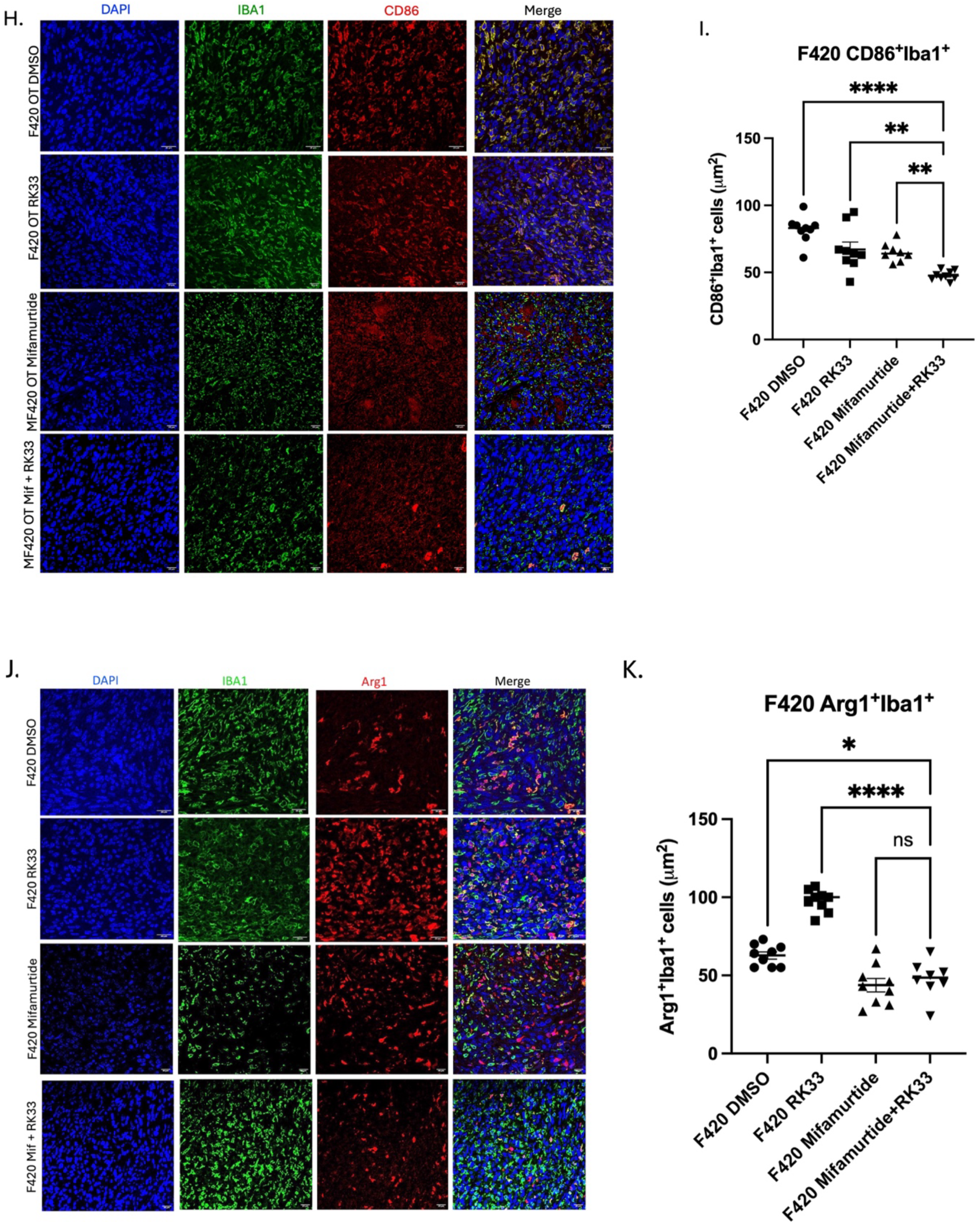
MTPE+RK33 prevents OS metastases. (A) Schematic of experimental design. (B) Mice were divided into four treatment groups (DMSO, RK33, Mifamurtide, RK-33+Mifamurtide) and treated for two weeks from time of tumor engraftment until amputation. Post-amputation, mice were treated for 8 more weeks until necropsied and assessed for metastasis. A statistically significant difference by Fisher’s exact test (p< 0.03) was seen in the number of mice with metastases in the combination treatment cohort (12 with metastases, 14 without metastases) compared to the number of mice with metastases in the control group (16 with metastases, 4 without metastases). (C) Representative photomicrographs of immunofluorescence analysis of macrophage polarization from F331 tumors treated with DMSO, RK33, Mifamurtide, RK-33+Mifamurtide. M1 phenotype: CD86+ (red) and Iba1+ (green) Mag=20*μm*. (D) Quantitative analysis of M1 macrophage phenotype (CD86^+^Iba1^+^) (p<0.0001). (E) Representative photomicrographs of immunofluorescence analysis of macrophage polarization from F331 tumors treated with DMSO, RK33, Mifamurtide, RK-33+Mifamurtide. M2 phenotype: Arg1+ (red) and Iba1+ (green). Mag=20*μm*. (F) Quantitative analysis of M2 macrophage phenotype (Arg1^+^Iba1^+^) (**** p<0.0001, * p=0.0221). (G) Mice were divided into four treatment groups (DMSO, RK33, Mifamurtide, RK-33+Mifamurtide) and treated for two weeks from time of tumor engraftment until amputation. Post-amputation mice were treated for 8 more weeks until necropsied and assessed for metastasis. No difference was seen in the number of mice with metastases seen in any of the cohorts. (H) Representative photographs of double immunostaining of macrophage polarization from MF420 tumors treated with DMSO, RK33, Mifamurtide, RK-33+Mifamurtide. M1 phenotype: CD86+ (red) and Iba1+ (green) Mag=20*μm*. (I) Quantitative analysis of M1 macrophage phenotype (CD86^+^Iba1^+^) (**** p<0.0001, ** RK-33 v. RK-33+Mifamurtide p=0.0011, ** Mifamurtide v. RK-33+Mifamurtide p=0.0052). (J) Representative photomicrographs of immunofluorescence analysis of macrophage polarization from F331 tumors treated with DMSO, RK33, Mifamurtide, RK-33+Mifamurtide. M2 phenotype: Arg1+ (red) and Iba1+ (green). Mag=20*μm*. (K) Quantitative Analysis of M2 macrophage phenotype (Arg1^+^Iba1^+^) (**** p<0.0001, * p=0.0115).

## Discussion

Our results demonstrate that DDX3 is a key modulator of interactions between bone sarcomas and the innate immune system. RK-33, a novel DDX3 inhibitor, is preferentially cytotoxic to sarcoma cells *in vitro* and inhibits growth of Ewing sarcoma xenografts expressing high DDX3 *in vivo*^17^. We found that both genetic and pharmacologic inhibition of DDX3 results in accumulation of cytoplasmic dsRNA in sarcoma cells, leading to a Type I interferon response. Sarcomas are largely unresponsive to immunotherapies due to an immunosuppressive tumor microenvironment. In addition to being directly toxic to sarcomas, by stimulating an innate immune response, we believe RK-33 has the potential to, in combination with other drugs, open new immunotherapeutic avenues to better treat patients with high-risk disease.

Inhibiting DDX3 in breast cancer cells also leads to an accumulation of dsRNAs^23^. Our results extend this finding to EWS and OS, demonstrating that DDX3 plays a role in regulating endogenous dsRNA levels across multiple malignancies. Mechanistically, dsRNAs are recognized by cytosolic RIG-I like receptors (RLRs) such as RIG-I and MDA5, which triggers a MAVS-dependent signaling cascade^39^. While RIG-1^40^ and MDA5^41^ exhibit distinct substrate preferences, recognizing short (10-40bp) and long (greater than 500bp) dsRNAs respectively, their collective activation culminates in a Type I interferon response and the upregulation of interferon stimulated genes. Interestingly, our DDX3 knockdown models exhibited elevated basal levels of dsRNA, suggesting that DDX3 is essential for maintaining normal RNA homeostasis. *In vitro*, RK-33-treated cells exhibited robust upregulation of downstream ISGs including IFI44L, OAS1/2, IFIT2/3, ISG15 and STAT1 in the absence of the induction of the upstream regulators IRF3, IRF5 and IRF7, suggesting that while dsRNA accumulation is sufficient to activate a downstream interferon response, tumor cells alone do not fully engage upstream innate sensing pathways. In contrast, RK-33-treated xenograft models demonstrate not only upregulation of ISGs but also robust upregulation of IRF3, IRF5 and IRF7, along with an expanded inflammatory cytokine and chemokine profile, including IL-6, CCL4 and CXCL10. These findings support a model in which cytoplasmic dsRNA generated by DDX3 inhibition is more effectively sensed *in vivo*, where host innate immune cells, potentially including macrophages, respond to dsRNA through Toll-like receptors^42,43^ and cytosolic RNA sensors^44,45^, leading to IRF-dependent interferon signaling. This may, in turn, enhance STAT1-mediated ISG expression in tumor cells via paracrine mechanisms. Collectively, these results suggest that DDX3 inhibition may not only induce a tumor-intrinsic antiviral-like state but also contribute to engagement of the innate immune microenvironment, resulting in a broader interferon associated inflammatory response *in vivo*. This interpretation is supported by gene set enrichment analysis (GSEA) of our RNA-seq data, which demonstrates strong enrichment of multiple inflammatory and cytokine signaling pathways. In contrast, the relatively modest changes observed in antigen presentation pathways suggest that response to DDX3 inhibition is predominantly driven by innate immune signaling, with limited evidence of robust activation of adaptive immune programs. The RNA sequencing data also help clarify the seemingly paradoxical downregulation of ISGs observed in F331 cells in the targeted qPCR panel. While interferon response genes remained downregulated in F331 tumors evaluated by RNA sequencing, consistent with the qPCR findings, pathway enrichment analysis revealed broader activation of inflammatory and cytokine signaling programs, suggesting that DDX3 inhibition in F331 engages pro-inflammatory transcriptomic programs that extend beyond the interferon axis captured by the targeted gene panel. The observation that distinct inflammatory pathways are activated across different model systems raises the question of what drives the heterogeneity in downstream signaling. The precise molecular makeup and genomic origins of the accumulating dsRNAs remains to be elucidated. Identifying the origin of these RNAs could better predict which downstream pathways are activated and broaden our understanding of the additional signaling events triggered by the specific types of dsRNA that accumulate in response to DDX3 inhibition.

To better understand how inhibition of DDX3 affects the tumor innate immune microenvironment, we characterized the macrophage population. Inhibition of DDX3 with RK-33 is a potent driver of myeloid reprogramming within the TME. Using a first-of-its-kind CyTOF 40+ macrophage marker panel, we observed a significant shift from pro-tumorigenic M2-like phenotype toward an anti-tumorigenic M1-like state in our ESX3 (Ewing sarcoma) and F331 (osteosarcoma) models. This shift is particularly striking within the M2c and M2d subpopulations, which are associated with immune suppression and tissue remodeling^46,47,48^. This transition likely stems from the upregulation of interferon stimulated genes observed following RK-33 treatment, creating an inflammatory milieu that inherently favors M1 polarization. By integrating CyTOF with immunofluorescence, we validated that these phenotypic shifts are reflected within the intact tumor architecture. The spatial data further demonstrates that the M1 rich environment induced by RK-33 is maintained across the entire tumor, indicating that macrophages are broadly distributed throughout the TME rather than limited to necrotic niches. This spatial distribution is critical, as effective cell-mediated cytotoxicity depends on the physical proximity of M1 macrophages to tumor cells^6^. It is notable that we observed a different phenotype in the F420 (osteosarcoma) model. In these tumors inhibition of DDX3 caused an enrichment of M2 macrophages, highlighting a critical intersection between oncogenic signaling and immune modulation. F420 is characterized by MYC overexpression, a known driver of metabolic reprogramming and immune evasion^49,50,51^. It is likely the MYC signaling in this model overrides the pro-inflammatory signaling triggered by DDX3 inhibition. In addition, MYC places exceptional metabolic and translational demands on the tumor cells, creating a state of heightened cellular stress. When DDX3 is inhibited in this context, the compounded photostatic stress may activate signals associated with cell damage and death, including efferocytosis cues, such as IL-10 and TGF-β that could direct macrophages toward a tissue-repair and clearance phenotype rather than an antitumor state^52,53,54^. Together, MYC-driven immunosuppressive signaling and stress-induced efferocytosis cues may create a microenvironment that is refractory to productive M1 polarization, potentially explaining why robust ISG upregulation in F420 tumors does not translate into an antitumor macrophage response. Whether this convergence directly underlies the M2 polarization observed in F420 remains to be proven but provides a mechanistic framework for understanding how the oncogenic landscape can render the innate immune microenvironment resistant to DDX3 inhibition.

Our study demonstrates that the combination of the DDX3 inhibitor RK-33 and the immunostimulant mifamurtide exerts a synergistic effect on reducing metastatic burden in osteosarcoma. While mifamurtide is currently clinically utilized in the EU to activate macrophages and monocytes^55,56^, our data suggests that its efficacy can be significantly enhanced when paired with DDX3 inhibition. In the F331 model, this combination led to a statistically significant reduction in metastatic lesions (p=0.03) compared to control. This therapeutic benefit is accompanied by a dramatic shift in the macrophage landscape characterized by a significant increase in pro-inflammatory CD86^+^Iba1^+^ macrophages and a concomitant decrease in immunosuppressive Arg1^+^Iba1^+^ macrophages. These findings suggest that RK-33 primes the TME by inducing a Type I interferon response, thereby creating a more receptive landscape for mifamurtide-induced macrophage activation. Critically, the F420 model showed a stark divergence in treatment response. Despite dual treatment, F420 tumors showed no reduction in metastatic growth (p=0.9), and exhibited a decrease in M1 macrophages with an increase in the M2 population. Our analysis suggests that MYC overexpression, the primary molecular difference between these models, acts as a dominant driver of this resistance^57^. The resistance observed in the MYC overexpressing model has significant implications for patient stratification in the clinic. Our results suggest that OS patients with high MYC expression levels may derive less benefit from the RK-33/mifamurtide combination, so clinical trials might be either limited to patients without MYC amplification or might be stratified based on MYC status. Future research should focus on whether the co-inhibition of MYC could unharness the therapeutic potential of RK-33 in currently resistant models. Characterizing the molecular crosstalk among DDX3, MYC, and macrophage polarization pathways will be essential to expanding the utility of these immunotherapeutic strategies across the heterogenous landscape of osteosarcoma.

In summary, we have shown that inhibition of DDX3 promotes a robust inflammatory environment and synergizes with mifamurtide to reduce metastatic burden (Figure 9). Mechanistically, DDX3 inhibition triggers dsRNA accumulation, driving the expression of innate immune signaling pathways and polarizing macrophages toward a pro-inflammatory M1 phenotype. By prioritizing the innate immune response over traditional checkpoint inhibition and other immunotherapy approaches focused on the adaptive immune system, this study identifies a novel therapeutic strategy. Since mifamurtide is already in clinical use, and a phase I study of RK-33 is anticipated to begin in 2026, the translation of RK-33 could rapidly establish a dual-agent regimen capable of improving survival for metastatic patients, who have faced a stagnant therapeutic landscape for decades.

**Figure 9:**
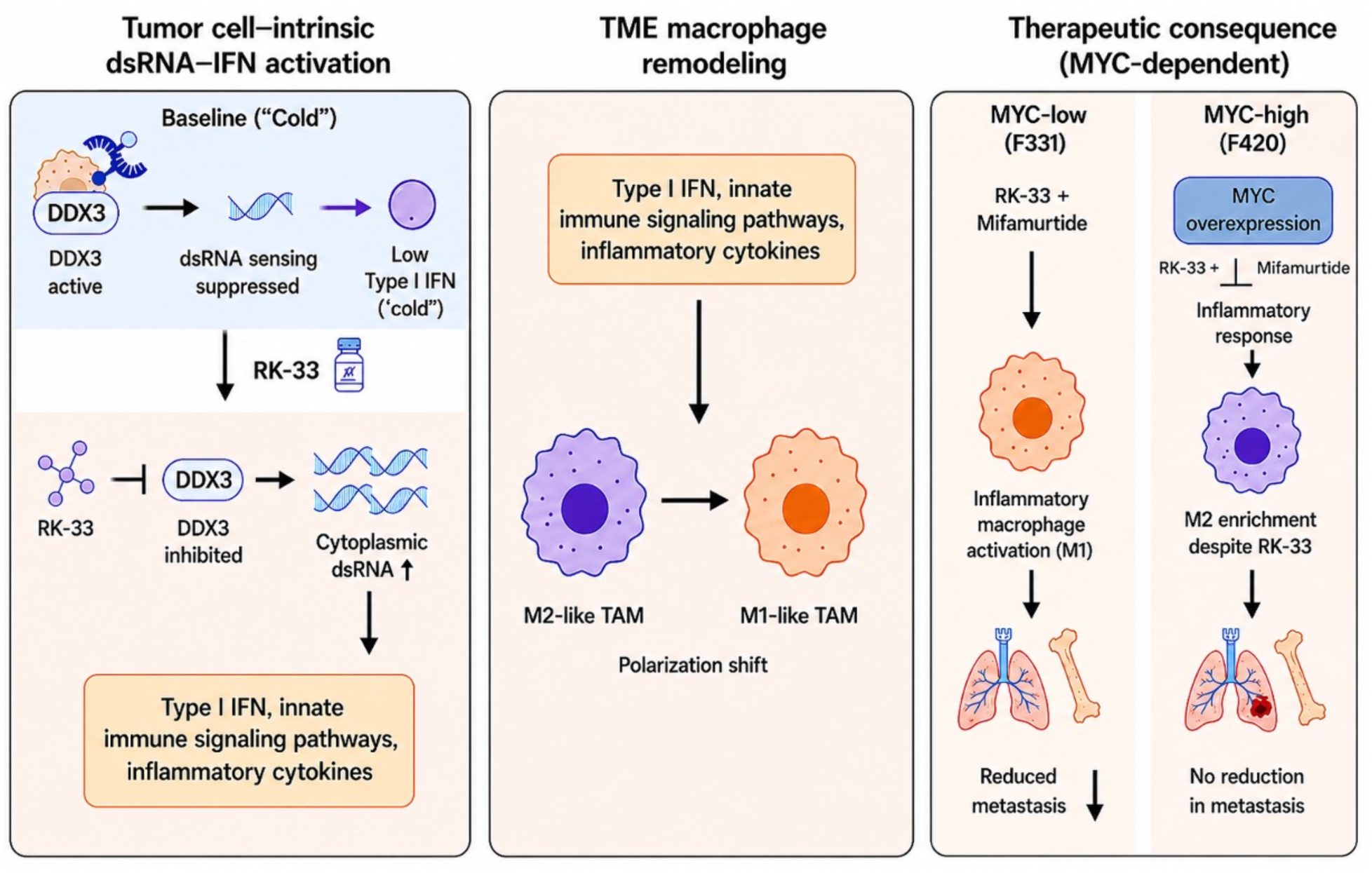
Proposed model of DDX3 inhibition; mediated innate immune activation, macrophage remodeling, and therapeutic response in bone sarcomas. When DDX3 is intact, dsRNAs are unwound, preventing the activation of a downstream inflammatory response. When DDX3 is inhibited by RK-33, it leads to an accumulation of dsRNAs triggering a downstream Type I IFN response. This Type I IFN response and cytokine release leads to a polarization shift from M2-like macrophages to M1-like. Together the innate immune activation and macrophage remodeling contribute to preventing metastasis in MYC-low tumors in response to RK-33 and mifamurtide.

## Materials and Methods

### Cell lines and xenografts

Established Ewing sarcoma cell lines TC-71 (RRID: CVCL_S882), TC-32 (RRID: CVCL_71571), A4573 (RRID: CVCL_6245) and established osteosarcoma cell lines HOS (RRID: CVCL_0312), SaOS2 (RRID: CVCL_0548), and U2OS (RRID: CVCL_0042) were originally purchased from ATCC and were cultured in 125 cm^2^ tissue culture flasks supplemented with RPMI-1640 (Gibco, Catalog #11875093) + 10% FBS and maintained under standard conditions. Cells were incubated in a ThermoScientific HERACell Vios 160i CO_2_ incubator at 37 °C. Cells were split every 3 days and incubated at different confluences per experiment. Cell line authentication was performed using STR analysis by the Genomics Core at the Albert Einstein College of Medicine. Periodic assessment for mycoplasma contamination was performed using Lonza Mycoalert Mycoplasma Detection kit. Cells were used for experiments within 3-5 passages of thawing. DDX3 shRNA knockdown cell lines were established as previously described^17^. We utilized two Ewing sarcoma patient-derived xenografts (PDXs) for *in vivo* experiments; JHHESX1 and JHHESX3. These EWS xenografts were generated in our laboratory after obtaining written informed consent under a tumor banking protocol approved by Johns Hopkins University School of Medicine IRB. For osteosarcoma, we used the DAR xenograft, created from cells isolated from a malignant pleural effusion in an osteosarcoma patient, gifted from Dr. Chand Khanna (NCI, NIH). All xenografts were grown in NOD/SCID/IL-2Rγ null (NSG) (RRID: IMSR JAX:005557) mice originally obtained from Jackson Laboratories and then bred by our group.

Osx-Cre, *Rb1*^lox/lox^, *Trp53*^lox/lox^ mice have been previously described^32,58^. First, Rb1^lox/lox^ mice were crossed with Trp53^lox/lox^ mice to generate Trp53^lox/lox^, Rb1^lox/lox^ mice, which were further crossed with Osx-Cre mice to generate Osx-Cre; Trp53^lox/lox^, Rb1^lox/lox^ (DKO). These mice were gifted by Dr. Bang Hoang (AECOM). Syngeneic murine cell lines^33^ F331 and F420 were gifted by Dr. Jason Yustein (Baylor College of Medicine). Both syngeneic mouse cell lines were grown in C57BL/6 mice from Jackson Laboratory. All mouse procedures were performed according to protocols approved by the Johns Hopkins Animal Care and Use Committee and the Institutional Animal Care and Use Committee at Albert Einstein College of Medicine.

### Tissue Microarray Immunohistochemistry

The OS tissue microarrays were provided by the Children’s Oncology Group through project ABTR14B2-Q. Following deparaffinization in xylene, the samples were rehydrated in decreasing ethanol dilutions. Endogenous peroxidase activity was blocked by endogenous peroxidase from Novolink Polymer Detection System (Leica Microsystems) and was followed by antigen retrieval by boiling for 20 minutes in EDTA buffer (pH 9.0). Slides were blocked with protein block from Novolink Polymer Detection System and subsequently incubated in a humidified chamber for 1 hour with anti-DDX3 (1:50, mAB AO196, RRID:AB_236197, Sigma Aldrich). Post primary block, secondary antibodies and diaminobenzidine treatment were performed with Novolink Polymer Detection System according to the manufacturer’s instructions. The slides were lightly counterstained with hematoxylin and mounted. Images were scored for staining intensity using a scale of 0 to +3 by Dr. Paul J. van Diest, University Medical Center Utrecht, The Netherlands. The slides were scanned using a Hamamatsu, NanoZoomer XR C1200-21/-22.

### Immunofluorescence

For immunofluorescent staining, cells were fixed with 4% paraformaldehyde for 15 minutes, washed with PBS, permeabilized using 0.2% Triton-X in PBS for 15 minutes, then blocked for 30 minutes using 5% goat serum and 1% bovine serum albumin (BSA) in 0.2% Triton-X PBS blocking buffer. Slides were then incubated with primary antibody J2 (Cell Signaling, Catalog #76651; 1:100) diluted in blocking buffer overnight at 4 °C. Slides were washed 2 times with 0.1% Tween 20 PBS the next day followed by 1 hour incubation at room temperature with fluorophore labeled secondary antibody AF488 goat-anti-mouse (ThermoFisher, Catalog # A-21131; 1:250) diluted in blocking buffer. Slides were then washed 3x with PBS and mounted with coverslips to slides using ProLong Gold Antifade Reagent with DAPI (Cell Signaling, Catalog #8961). Slides were imaged using AECOM’s Analytical Imaging Facility’s Leica SP5 confocal microscope at 40x and 63x magnification. Images were processed and quantified using Volocity software (RRID: SCR_002668, Quorum Technologies). Tumor sample slides were first deparaffinized and rehydrated through xylene, graded alcohols to distilled water, followed by antigen retrieval by boiling in a sodium citrate buffer (pH 6). The slides were then permeabilized using 0.4% Triton-X in PBS with 1% goat serum, followed by blocking for 30 minutes in 0.1% Triton-X in PBS with 5% goat serum. Slides were then incubated overnight in primary antibody (CD86; BD Bioscience, Catalog # 553689; 1:100) (Arg1; GeneTex, Catalog # GTX113131; 1:500) (IBA1; Invitrogen, Catalog # MA5-27726; 1:500) diluted in blocking buffer. The next day slides were washed twice with 1% goat serum, 0.1% Triton-X in PBS and then incubated for an hour at room temperature with fluorophore labeled secondary antibodies (AF647 Goat anti Rat; ThermoFisher, Catalog # A-21241; 1:250) (AF555 Goat anti Rabbit; ThermoFisher, Catalog # A-21429; 1:250) (AF488 Goat anti Mouse; ThermoFisher, Catalog # A-11001; 1:250) diluted in blocking buffer. Slides were then washed 3x with PBS and mounted the coverslips to slides using ProLong Gold Antifade Reagent with DAPI. Slides were imaged using AECOM’s Analytical Imaging Facility’s Leica Stellaris 8 confocal microscope at 40x and 63x magnification. Images were processed and quantified using ImageJ software (RRID: SCR_003070, NIH).

### RNA Isolation and qRT-PCR

RNA was isolated from cell lines and mouse tumors using the RNeasy kit (Qiagen Catalog #74104) and reverse-transcribed into cDNA using the Iscript SuperScript reverse transcriptase RT-PCR kit (Bio-Rad, Catalog #1708840). For all cell lines, 300,000 cells were plated per well in a 6-well, tissue culture-treated plate (Corning, Catalog #3516) with 3 mL of RPMI-1640 (Gibco, Catalog #11875093) + 10% FBS and maintained under standard conditions. After 24 hours to allow for adherence, the cells were treated with 2 µM of RK-33 or an equivalent amount of DMSO. Mouse tumors were stored in RNAlater (Invitrogen, Catalog #AM7021) until RNA was extracted following the same method. All primers were obtained from Bio-Rad (Supplemental Table 1) and used with SYBR Green master mix (Bio-rad, Catalog #1725272). Primers used for MYC assessment were obtained from OriGene (Catalog #MP208493). Quantitative RT-PCR was performed using the Applied Biosystems QuantStudio5 Real-Time PCR thermal cycler. The cycle threshold (Ct) values were used to calculate fold change in expression using the 2^− ΔΔCt^ method, and qRT-PCR and melting curves analysis were conducted using the ThermoFisher data analysis cloud. Gene expression heatmap for PCR arrays was created using the Morpheus software (https://software.broadinstitute.org/morpheus/).

### RNA Sequencing

RNA was isolated using the Qiagen RNeasy kit (Qiagen Catalog #74104). Libraries were constructed and sequenced by the Yale Center for Genome Analysis (YCGA) and by Plasmidsaurus.

### Differential Gene Expression Analysis (F331 v. F420)

Quality of the fastq files was assessed using FastQC v0.12.1. Reads were then quality filtered using fastp v0.24.0 with poly-X tail trimming, 3’ quality-based trail trimming, a minimum Phred quality score of 15, and a minimum length requirement of 50 bp. Quality-filtered reads were aligned to the reference genome using STAR aligner v2.7.11 with non-canonical splice junction removal and output of unmapped reads, followed by coordinate sorting using samtools v1.22.1. PCR and optical duplicated were removed using UMI-based deduplication with UMIcollapse v1.1.0. Alignment quality metrics, strand specificity, and read distribution across genomic features were assessed using RSeQC v5.0.4 and Qualimap v2.3, with results aggregated into a comprehensive quality control report using MultiQC v1.32. Gene-level expression quantification was performed using featureCounts (subread package v2.1.1) with strand-specific counting, multi-mapping read fractional assignment, exons and three prime UTR as the feature identifiers, and grouped by gene_id. Final gene counts were annotated with gene biotype and other metadata extracted from the reference GTF file. Differential expression was done with edgeR v4.0.16 using standard practice including filtering for low-expressed genes with edgeR::filterbyExpr with default values. Functional enrichment, when available, is performed using gene set enrichment analysis with gseapy v0.12 using MSigDB Hallmark gene set.

### Differential Gene Expression Analysis (RK-33 vs DMSO)

Raw fastq files for PDXs (10 DMSO and 12 RK-33 treated) and GEMMs (6 DMSO and 7 RK-33 treated) were processed to obtain transcript-level expression estimates using Salmon^59^ (version 1.10.3). The human and mouse reference transcriptomes were obtained from GENCODE^55^ release v45 (GRCh38.p14) and vM34 (GRCm39) respectively. DESeq2^60^ (version 1.34.0) was used to perform differential expression analysis with default parameters. Prior to differential gene expression analysis, the lowly expressed genes were filtered out. Specifically, only genes that achieved a minimum of 10 counts in at least 6 samples in the GEMM comparison and at least 10 samples in the PDX comparison were retained. Enrichment analysis was performed using enrichR^61^ (version 3.2) using GO_Biological_Process and KEGG gene sets.

### Mass Cytometry (CyTOF) Analysis of Mouse Tumor Macrophages

#### CyTOF Antibody Panel Design and Metal Conjugation

A macrophage focused CyTOF panel was designed using the Standard BioTools panel designer platform (https://standardbio.com). Antibodies were obtained either pre-conjugated from Standard Biotools (SBT) or conjugated in-house using either the Maxpar® X8 or MCP9 Antibody Labeling Kits (SBT https://standardbio.com/support/instrument-support/helios-and-cytof-2-support/) according to the manufacturer’s instructions. Each antibody was titrated using a Universal 10-Tube Titration for Mass Cytometry protocol^62^ to ensure optimal signal resolution and minimal background staining.

To facilitate antibody titration, we used cells that closely resembled the experimental tumor-derived macrophages, ensuring minimal technical variability and allowing reliable comparison of staining intensities across antibody concentrations. Prior to performing titration experiments, *in vitro* polarization of macrophage subtypes was carried out following the protocol described by Xin et al^63^. The complete list of surface and intracellular markers can be found in Supplemental Table 2.

#### Tumor Tissue Harvest and Preparation

Primary tumors were dissected aseptically and immediately placed into MACS Tissue Storage Solution (Miltenyi Biotec, Catalog # 130-100-008) to preserve cell viability. Samples were kept refrigerated and were processed within 48 hours of collection. Tumor tissues were dissociated into single cell suspensions following STAR Protocol^64,65^. After enzymatic and mechanical dissociation, cell suspensions were filtered, washed in DPBS, and resuspended in freezing medium (90% FBS + 10% DMSO). Cells were aliquoted into cryovials and frozen overnight in Styrofoam at -80 °C, then stored long term in the vapor phase of liquid nitrogen. On the day of CyTOF staining, cell suspensions were thawed in a 37°C water bath and transferred into prewarmed complete RPMI –1640 (Gibco) + 10% FBS (CRPMI). Cells were activated for 5 hours with PMA (25 ng/mL; Phorbol 12-myristate 13-acetate), Ionomycin (1 µg/mL), and Brefeldin A (5 µg/mL) in CRPMI. This activation step was performed to re-initiate cytokine production and prevent secretion prior to intracellular staining (adapted from Lin et al and optimized for our sample sets and macrophage populations^66^).

Following activation, cells were washed twice by adding 2mL of Maxpar® Cell Staining Buffer (SBT, Catalog #201068) and centrifuged at 300xg for 5 minutes. Prior to surface antigen staining live/dead discrimination was performed using Cell-ID™ Cisplatin (198Pt) (SBT, Catalog #201064), to label membrane-compromised cells and enable exclusion of dead cells during downstream gating. After viability staining, cells were washed twice in Maxpar® Cell Staining Buffer before proceeding to staining using the Maxpar® Cytoplasmic or Secreted Antigen Staining with Fresh Fix protocol, following the manufacturer’s recommended workflow (https://content.ilabsolutions.com/wpcontent/uploads/2019/06/Cytoplasmic_Secreted_Antigen_Staining_Protocol.pdf). Surface antigen staining was performed with the antibodies listed (Supplemental Table 2). Cells were then fixed overnight in 2% paraformaldehyde (PFA) (Life Technologies, Catalog # 28908) at 4 °C. After fixation, cells were washed and stained with intracellular antibodies in permeabilization buffer for 30 minutes. (Ebioscience Permeabilization Buffer, Catalog # 00-8333-56). After the intracellular staining, cells were washed twice with Maxpar Cell staining buffer and then fixed in 1.6% PFA for 10 minutes, followed by overnight fixation at 4 °C in Maxpar Fix and Perm buffer (SBT – Catalog # 201067) containing Cell-ID™ Intercalator-Ir for DNA labeling according to established protocols (SBT; Cell-ID Intercalator-Ir, Cat# 201192). The following day, cells were washed thoroughly with Maxpar Cell staining buffer followed by Maxpar Cell Acquisition Solution (CAS) (SBT, Catalog # 201240) to remove residual fixative and ensure proper sample preparation for acquisition on the CyTOF instrument. Before acquisition, cells were resuspended at 0.5–1 × 10⁶ cells/mL in CAS containing EQ™ Four-Element Calibration Beads (SBT, catalog #201078) for normalization.

#### CyTOF Data Normalization and Analysis

Acquired FCS files were normalized using bead-based normalization in the instrument software (Helios/CyTOF Software v7.1). The resulting .fcs files were processed in OMIQ (www.omiq.ai), where the data was gated for live singlets and exclusion of highly cisplatin-positive dead cells. High-dimensional analysis was performed using UMAP (neighbors = 15, minimum distance = 0.4, Euclidean metric) and Phenograph clustering (k = 20, Leiden method).

#### Protein Extraction and Western Blot Analysis

Total cell extracts were prepared using RIPA buffer with protease inhibitors (SantaCruz, Catalog #sc-24984A) with additional PMSF (1mM), sodium orthovanadate (1mM) and sodium fluoride (50mM) added immediately preceding use. Protein concentration was quantified using the Pierce BCA colorimetric assay (ThermoFisher, Catalog #23225) and quantified using BSA standards with a Multiskan plate imager at 750nM. Extracted proteins were run on 4-15% TGX gels (Bio-Rad Catalog #456108) and transferred onto methanol-activated PVDF membranes using BioRad dry transfer system. Immunoblots were developed with c-MYC (Cell Signaling, Catalog #18583; 1:1000) and HSP90 (Cell Signaling, Catalog #4874; 1:1000) primary antibodies and corresponding anti-rabbit-HRP (Vector labs, Catalog #MP7451; 1:10,000) secondary antibody. Blots were scanned using the ThermoFisher myECL imager.

#### *In vivo* Studies

Tumor fragments approximately 5-6 mm^3^ in volume were coated with Matrigel (BD Biosciences) and surgically implanted into the tibia of experimental mice. All mice at time of implant were between 6-8 weeks old and to ensure that the sex of the mouse would not bias results, each experiment contained equal numbers of males and females. Once tumors were palpable at around 7 mm in diameter, mice were randomly divided into 4 cohorts depending on treatment received: control (DMSO only), RK-33 only (from the Raman laboratory, Johns Hopkins University), mifamurtide only (MedChemExpress, Catalog #HY-13682), or RK-33 + mifamurtide. Either 10 mg/kg RK-33 or equivalent 50 µl DMSO was injected intraperitoneally 5 days a week until the conclusion of the experiment (about 2 weeks of tumor growth to amputation and about 8 weeks post amputation). Mifamurtide (1 mg/kg) was injected retro-orbitally twice a week for the same time course. Tumor dimensions were measured twice weekly using calipers until reaching a diameter of 15 mm. Once the tumors reached this diameter, the limbs were amputated and the mice were followed for recurrence or metastasis. To evaluate for metastases, mice were sacrificed eight weeks after hindlimb amputation. Necropsy was performed and visible metastatic lesions were quantified and collected.

#### Statistical Analysis

All statistical analyses were conducted using GraphPad Prism 9 (GraphPad Software). T-test was used for data comparison between two groups, and analysis of variance (ANOVA) was used for data comparison between three or more groups. A Chi square, Fischer’s exact test was used for in vivo metastases analysis. A p<0.05 was considered statistically significant.

## Author Contributions

RW contributed to coordinating and designing the study; developing, acquiring, analyzing, and interpreting the data; and drafting the manuscript. PG, KQM, KW, KHS, JH, and JO contributed to the acquisition, analysis, and interpretation of data and reviewed and edited the manuscript. DP and PC contributed to analysis and interpretation of the data and reviewed and edited the manuscript. NTH, PJD contributed to acquisition and analysis of the data. VR and EUAS participated in reviewing and editing the manuscript. DML participated in the coordination and design of the study, interpretation of the data, and revision and review of the manuscript. All authors read and approved the final manuscript.

## Acknowledgments

We are grateful for the support of H. Guzik and V. DesMarais of AECOM Analytical Imaging Facility for training and technical assistance, supported by NCI cancer center support grant P30CA013330. This work utilized an advanced confocal microscope, the Leica Stellaris 8 that was purchased with funding from a National Institute of Health SIG grant 1S10OD023591-01. This work was funded by Alex’s Lemonade Stand Foundation and by the Montefiore Einstein Cancer Center Support Grant 2P30CA013330. Additional support for research was provided by a grant from the WWWW (QuadW) Foundation, Inc. (https://www.quadw.org/) to the Children’s Oncology Group, the COG Biospecimen Bank Grant U24CA196173, and the NCTN Operations Center Grant U10CA180886 to the Children’s Oncology Group from the National Cancer Institute of the National Institutes of Health.

