## Supplementary Figures for "DDX3 Regulates the Innate Immune Response to Bone Sarcomas"

Supplementary Figure 1: S1

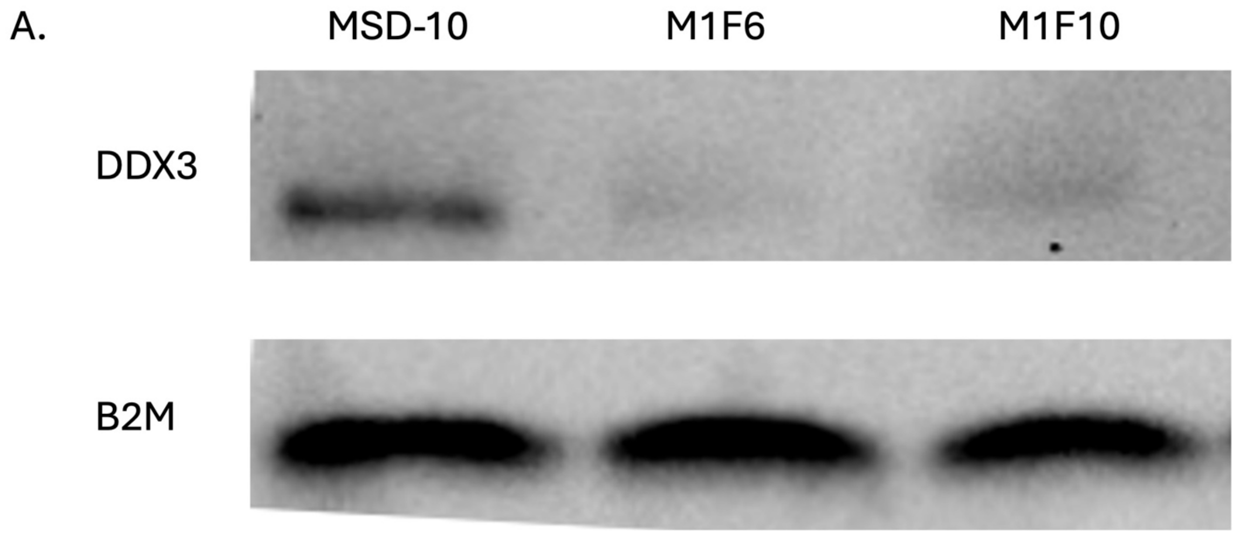


(A) Western blot confirming DDX3 knockdown expression in M1F6 and M1F10 cell lines compared to the scramble control (MSD10).

Supplementary Figure 2. S2

A.

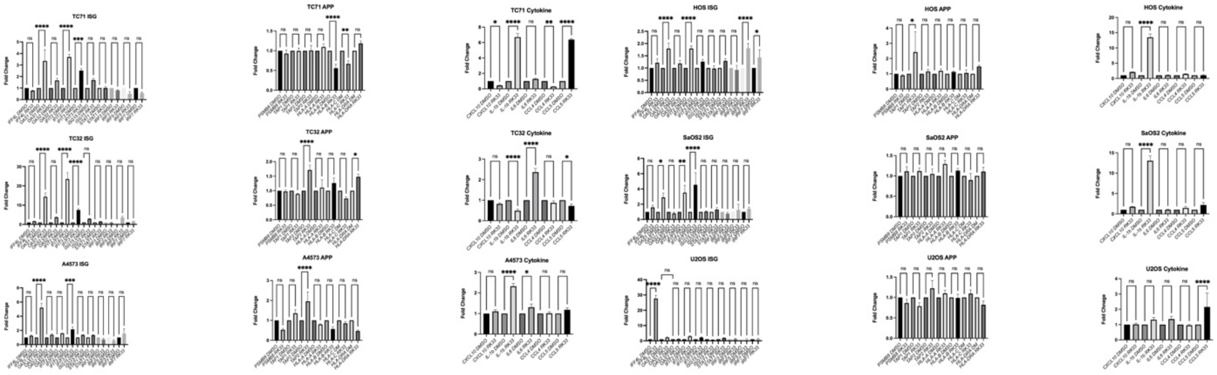


B.

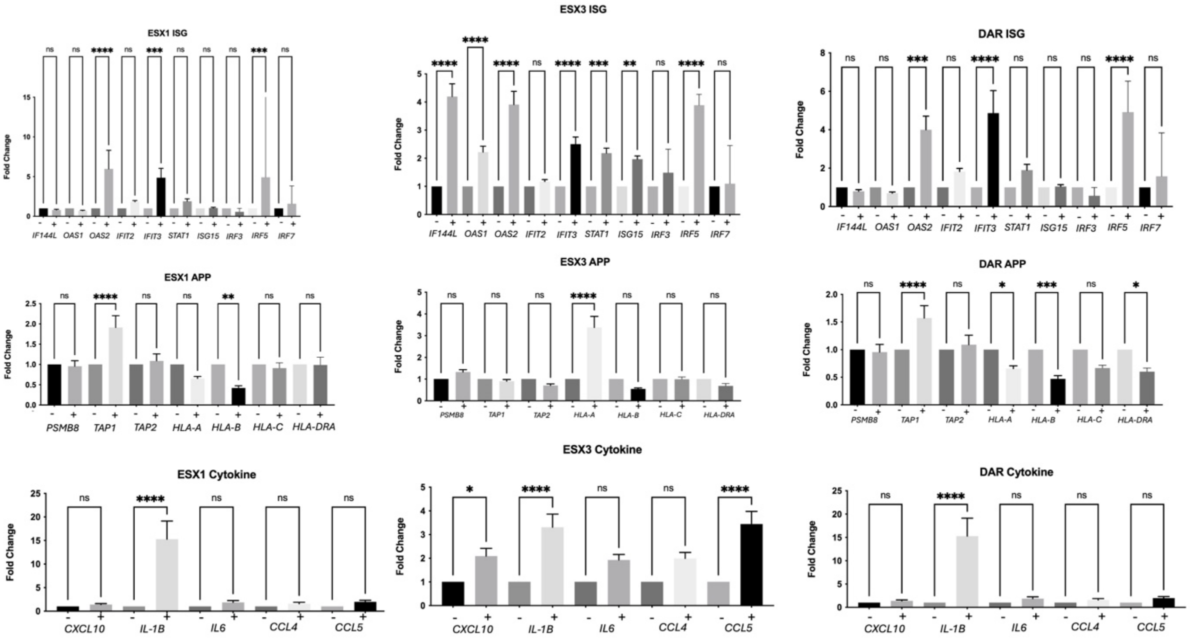

C.

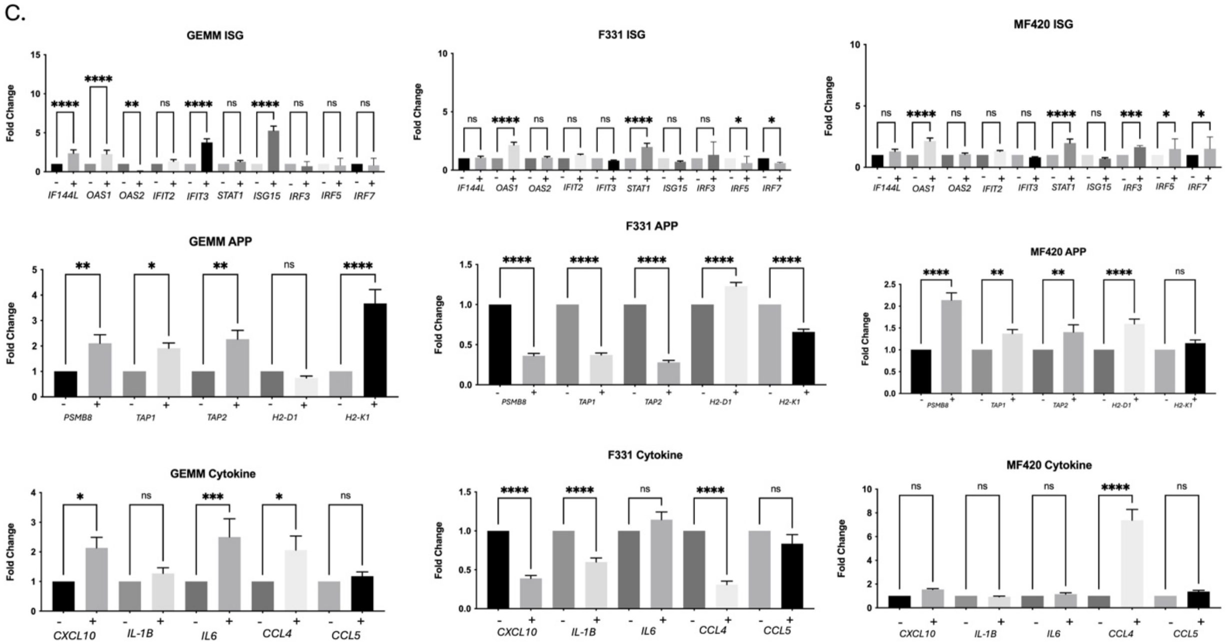


(A) qRT-PCR of mRNA expression from Ewing sarcoma cell lines (TC71, TC32, and A4573) and osteosarcoma cell lines (HOS, SaOS2, and U2OS) of ISGs, APPs, and cytokine and chemokines. Error bars are the SEM of triplicate experiments, and each experiment was repeated three times. (B) qRT-PCR of mRNA expression from Ewing sarcoma PDX tumors (ESX1 and ESX3) and Osteosarcoma PDX tumor (DAR) of ISGs, APPs, and cytokine and chemokines. Error bars are the SEM of triplicate experiments, and each experiment was repeated three times.(C) qRT-PCR of mRNA expression from immune competent models of osteosarcoma (GEMM, F331, and F420) of ISGs, APPs, and cytokine and chemokines. Error bars are the SEM of triplicate experiments, and each experiment was repeated three times.

Supplementary Figure 3: S3

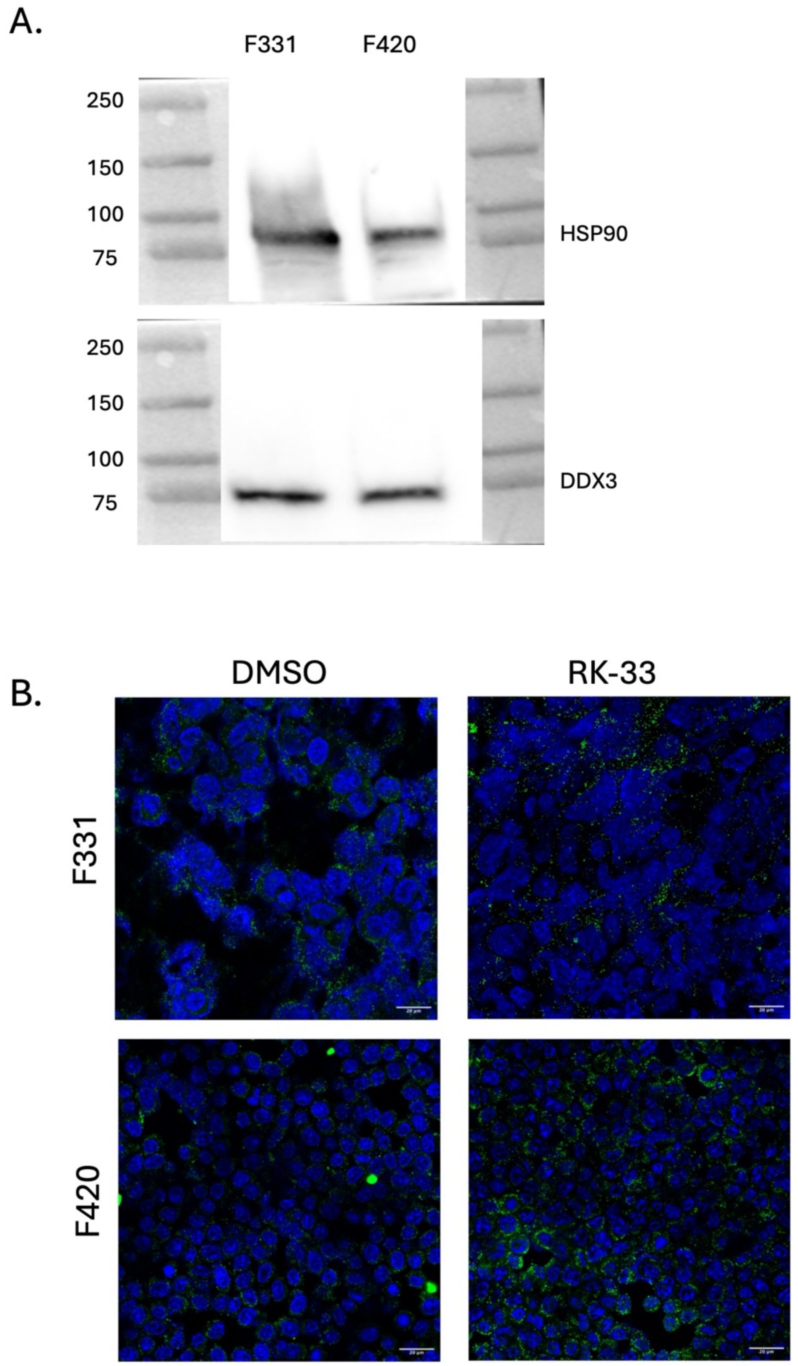


(A) Western blot confirming DDX3 expression in F331 and F420 cell lines. (B)Osteosarcome F331 and F420 cell lines analyzed by immunofluorescence following 24-hour treatment with either RK-33 or DMSO for accumulation of dsRNA with anti-dsRNA specific antibody (J2). Mag=20$\mu m$.
